# Regulation of Bud Emergence by a MAPK Pathway

**DOI:** 10.1101/786426

**Authors:** Aditi Prabhakar, Jacky Chow, Alan J. Siegel, Paul J. Cullen

**Author notes:** Corresponding author: Paul J. Cullen, Department of Biological Sciences, 532 Cooke Hall, State University of New York at Buffalo, Buffalo, NY 14260-1300. Author contributions: A.P. designed experiments, generated data, and wrote the paper; J.C. performed experiments and analyzed the data; A.J.S. performed experiments; P.J.C. designed experiments and wrote the paper. The authors have no competing interests in the study.

## Abstract

All cells establish and maintain an axis of polarity that is critical for cell shape and progression through the cell cycle. A well-studied example of polarity establishment is bud emergence in yeast, where the Rho GTPase Cdc42p regulates symmetry breaking at bud sites and the establishment of polarity by interacting with effector proteins. The prevailing view of bud emergence does not account for regulation by extrinsic cues or signal transduction pathways. Here, we show that the MAPK pathway that controls filamentous growth (fMAPK pathway), which also requires Cdc42p and the effector p21 activated kinase (PAK) Ste20p, regulates bud emergence under nutrient-limiting conditions that favor filamentous/invasive growth. The fMAPK pathway regulated the expression of polarity targets that included the gene encoding a direct effector of Cdc42p, Gic2p. The fMAPK pathway also stimulated GTP-Cdc42p levels, which is a critical determinant of polarity establishment. The fMAPK pathway activity was spatially restricted to bud sites and highest at a period in the cell cycle that coincided with bud emergence. Time-lapse fluorescence microscopy showed that the fMAPK pathway stimulated the rate of bud emergence during filamentous growth. Unregulated activation of the fMAPK pathway induced growth at multiple sites that resulted from multiple rounds of symmetry breaking inside the growing bud. Collectively, our findings identify a new regulatory aspect of bud emergence that sensitizes this essential cellular process to external cues.

## INTRODUCTION

All cells establish an axis of polarity, which is critical for cell shape, the organization of cellular compartments, and progression through the cell cycle (Doerr and Ragkousi, 2019). Cell polarity can be reorganized in response to intrinsic and extrinsic cues and is required during development and for other processes that require dynamic changes in cell shape such as cell migration and differentiation (Henderson *et al.*, 2018; Pei *et al.*, 2019; Piroli *et al.*, 2019). Defects in cell polarity are commonly associated with human disease. For example, the mis-regulation of cell polarity leads to metastasis in many types of cancers (Noguchi *et al.*, 2018; Taciak *et al.*, 2018; Fomicheva *et al.*, 2019).

Cdc42p and other members of the Rho family of GTPases are central regulators of cell polarity (Pringle *et al.*, 1995; Pruyne and Bretscher, 2000a, b; Irazoqui and Lew, 2004; Arkowitz and Iglesias, 2008; Zihni and Terry, 2015; Jimeno and Santos, 2017; Woods and Lew, 2019). In yeast, Cdc42p regulates bud emergence, which is an essential process where daughter cells or buds are produced from mother cells. The pathway that regulates bud emergence has been extensively studied and represents one of the best examples for how cells establish an axis of polarity (**Fig. 1A**, green pathway). Activation of Cdc42p by the guanine nucleotide exchange factor (GEF) Cdc24p at bud sites – or at random sites in bud-site-selection mutants – results in symmetry breaking, which commits the cell to grow at a particular site. Positive and negative feedback loops promote polarity establishment by amplifying the levels of active or GTP-bound Cdc42p at the incipient bud site (Irazoqui *et al.*, 2003; Kozubowski *et al.*, 2008; Chiou *et al.*, 2018; Woods and Lew, 2019). The GTP-bound conformation of Cdc42p binds effector proteins (Wu and Jiang, 2005; Wu *et al.*, 2006; Yang *et al.*, 2007; Guo *et al.*, 2010; Bendezu and Martin, 2013; McCormack *et al.*, 2013), including Gic1p and Gic2p (Brown *et al.*, 1997; Chen *et al.*, 1997; Kawasaki *et al.*, 2003; Iwase *et al.*, 2006; Liu *et al.*, 2019), p21 activated kinases (PAKs) Ste20p and Cla4p (Cvrckova *et al.*, 1995; Simon *et al.*, 1995; Gulli *et al.*, 2000; Hofmann *et al.*, 2004; Takahashi and Pryciak, 2007), and the formin Bni1p (Evangelista *et al.*, 1997; Sherer *et al.*, 2018). In a highly choreographed manner that is coordinated with the cell cycle (Moran *et al.*, 2019), effector proteins regulate the assembly of septin ring, which forms a diffusion barrier between mother and daughter cells, and the formation of actin cables, which direct the delivery of vesicles to the nascent bud site. In this way, buds are produced from mother cells in a precisely spatially and temporally regulated manner.

**Figure 1.**
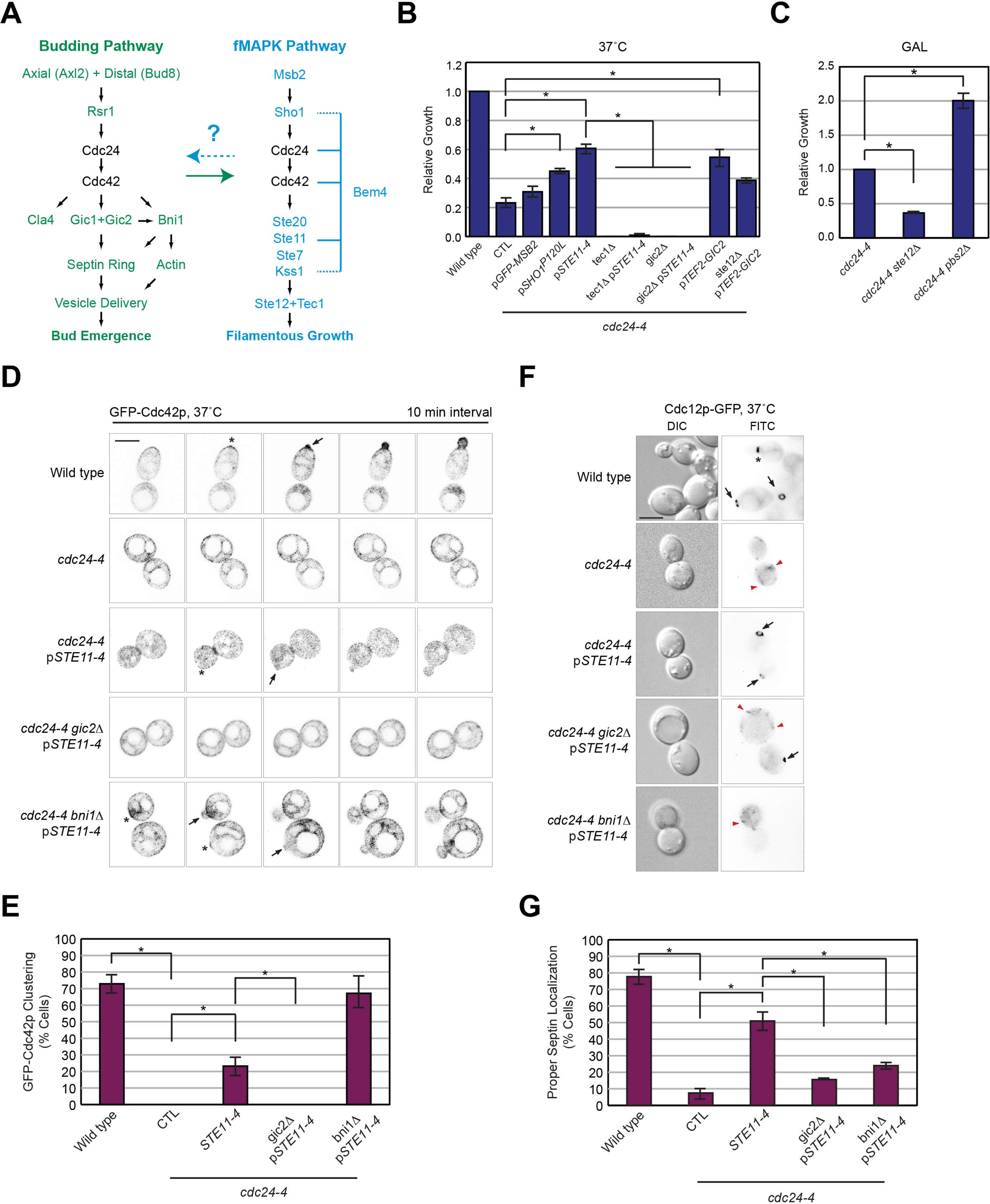
Role of the fMAPK pathway in rescue of the growth defect of the cdc24-4 mutant. **(A)** Pathways that regulate bud emergence (green) and filamentous growth (blue). Cdc24p and Cdc42p regulate both pathways. Not all proteins are shown. **(B)** Growth of the indicated strains relative to wild-type cells. p*GFP-MSB2*, p*SHO1*^P120L^, p*STE11-4* and p*TEF2*-*GIC2* were expressed from plasmids (see Table 2). Error bars show the standard error of the mean from three separate trials. Asterisk, p-value < 0.01. CTL, plasmid pRS316. **(C)** Growth of strains relative to *cdc24-4*. This experiment was performed in galactose (GAL) media, which also compromised the viability of the *cdc24-4* mutant, because the *pbs2*Δ mutant has a growth defect at 37°C (Winkler *et al.*, 2002). (**D**) Inverted maximum intensity projection of GFP-Cdc42p localization in the indicated strains, examined after incubation at 37°C for 4 h. Asterisks point to GFP-Cdc42p clustering. Arrows, sites of bud emergence. Bar, 5 microns. (**E**) Quantification of GFP-Cdc42p clustering. Error bars represent standard error of mean for three separate trials. 50 cells were counted in each trial. Asterisk, p-value < 0.05. (**F**) Cdc12p-GFP localization in the indicated strains, examined after incubation at 37°C for 4 h. Arrows, Cdc12p-GFP localization in incipient buds; asterisk, Cdc12p-GFP localization at mother-bud neck in growing bud; arrowheads, mislocalized Cdc12p-GFP. Bar, 5 microns. (**G**) Quantification of septin localization. Error bars represent standard error of mean for three separate trials. 50 cells were counted in each trial. Asterisk, p-value < 0.05.

Cdc42p also regulates <u>m</u>itogen <u>a</u>ctivated <u>p</u>rotein <u>k</u>inase (MAPK) pathways (Simon *et al.*, 1995; Brown *et al.*, 1996; Martin, 2019). MAPK pathways are evolutionarily conserved signaling modules that control cell differentiation and the response to stress in eukaryotes (Yoon and Seger, 2006; Raman *et al.*, 2007; Dinsmore and Soriano, 2018; Papa *et al.*, 2019). In yeast, nutrient limitation induces a Cdc42p-dependent MAPK pathway that regulates filamentous/invasive/pseudohyphal growth [**Fig. 1A**, blue pathway, fMAPK pathway (Gimeno *et al.*, 1992; Roberts and Fink, 1994; Mosch *et al.*, 1996; Peter *et al.*, 1996; Leberer *et al.*, 1997; Mosch *et al.*, 1999; Pan *et al.*, 2000; Gancedo, 2001; Borneman *et al.*, 2007; Adhikari *et al.*, 2015; Cullen, 2015). Filamentous growth occurs in many fungal species, and in some plant and animal pathogens, filamentous growth is required for virulence (Lo *et al.*, 1997; Desai *et al.*, 2014; Lagree and Mitchell, 2017). At the head of the fMAPK pathway, plasma-membrane proteins Msb2p and Sho1p regulate the Cdc42p module, which leads to the recruitment and activation of Ste20p, the specific Cdc42p effector that regulates the fMAPK pathway. Ste20p in turn induces a MAPK cascade composed of Ste11p (MAPKKK), Ste7p (MAPKK), and Kss1p (MAPK) (Roberts and Fink, 1994; Madhani *et al.*, 1997). The MAPK Kss1p regulates the activity of transcription factors Ste12p and Tec1p, as well as transcriptional repressors and other factors (Madhani and Fink, 1997; Bardwell *et al.*, 1998; van der Felden *et al.*, 2014) to regulate expression of target genes that bring about the filamentous cell type (Rupp *et al.*, 1999; Roberts *et al.*, 2000).

Despite the fact that bud emergence is extensively regulated by positive and negative feedback loops and is coordinated with the cell cycle, bud emergence is not known to be regulated by extrinsic cues or signaling pathways. The fMAPK and polarity pathways both require Cdc42p, which suggests that functional cross talk may occur between the two pathways. In support of this possibility, we have recently found that bud-site-selection proteins regulate the fMAPK pathway [**Fig. 1A**, green arrow, (Basu *et al.*, 2016)]. We tested the hypothesis that reciprocal cross talk might occur from the fMAPK pathway to the bud emergence pathway (**Fig. 1**, blue arrow question mark), and discovered a role for the fMAPK pathway in regulating bud emergence. An active fMAPK pathway rescued the bud emergence defect of a polarity mutant (*cdc24-4*), and was required for proper bud emergence during filamentous growth. The fMAPK pathway accomplished this role by inducing expression of a direct effector of Cdc42p that functions in budding (*GIC2*) and by directly stimulating GTP-Cdc42p levels. The activity of fMAPK pathway was regulated during the cell cycle to peak prior to and during bud emergence and when mis-regulated induced growth at multiple sites. Our study therefore identifies a new regulatory facet of bud emergence that is subject to regulation by a MAPK pathway and external cues. Given that Rho GTPases are functionally connected to MAPK pathways in higher eukaryotes, it is likely that polarity establishment is more likely to be subject to input from MAPK and other pathways than is currently appreciated.

## RESULTS

### fMAPK Pathway Rescues the Bud Emergence Defect of cdc24-4 Through Gic2p

*MSB2* (*<u>M</u>ulticopy <u>S</u>uppressor of <u>B</u>udding Defect 2*) was initially characterized as a high-copy suppressor of the budding and growth defects of the *cdc24-4* mutant (Bender and Pringle, 1989, 1992). Msb2p was subsequently identified as the mucin-type glycoprotein that regulates the fMAPK pathway [**Fig. 1A**, blue pathway (Cullen *et al.*, 2004)]. To determine whether Msb2p regulates budding through the fMAPK pathway, plasmids carrying alleles that hyper-activate the fMAPK pathway were introduced into the *cdc24-4* mutant and examined for growth at 37°C. Hyperactive versions of Msb2p (p*GFP*-*MSB2*) weakly suppressed the growth defect of the *cdc24-4* mutant (**Fig. 1B**). Hyperactive versions of Sho1p (p*SHO1*^P120L^) and Ste11p (p*STE11-4*) more robustly rescued the growth defect of the *cdc24-4* mutant (**Fig. 1B**). p*STE11-4* also rescued the polarity defect of the *cdc24-4* mutant (*Fig. S1A*). Therefore, activated versions of MAPK pathway components can rescue the growth and polarity defects of a bud emergence mutant.

Msb2p, Sho1p, and Ste11p regulate two MAPK pathways [fMAPK and HOG (Saito, 2010)]. A pathway-specific regulator of the HOG pathway, Pbs2p (Brewster *et al.*, 1993; Zarrinpar *et al.*, 2004; Nishimura *et al.*, 2016), was not required for growth of the *cdc24-4* mutant (**Fig. 1C**). In fact, the *cdc24-4*Δ *pbs2*Δ double mutant grew better than the *cdc24-4* single mutant, perhaps because the HOG pathway negatively regulates the fMAPK pathway and its loss results in hyperactive fMAPK (Davenport *et al.*, 1999). By comparison, a pathway-specific regulator of the fMAPK pathway, Tec1p (Madhani and Fink, 1997) was required for viability of the *cdc24-4* mutant (**Fig. 1B**). Tec1p functions with another transcription factor, Ste12p (Liu *et al.*, 1993), which was also required for growth of *cdc24-4* (**Fig. 1C**). Ste11p and Ste12p also regulate the mating pathway, but these experiments were carried out in a mating defective strain (*ste4*Δ). Tec1p was also required for suppression of the growth defect of the *cdc24-4* mutant by p*STE11-4* (**Fig. 1B**). p*STE11-4* also induced phosphorylation of the MAP kinase for the fMAPK pathway, Kss1p (P∼Kss1p) in the *cdc24-4* mutant (*Fig. S1B*). Therefore, the MAPK pathway that controls filamentous/invasive growth regulates bud emergence in a defined genetic context.

The fMAPK pathway controls filamentous growth by regulating target genes that control cell adhesion [*FLO11* (Rupp *et al.*, 1999)], bud-site selection [*BUD8* (Taheri *et al.*, 2000; Harkins *et al.*, 2001; Cullen and Sprague, 2002)] and cell elongation [*CLN1* (Kron *et al.*, 1994; Loeb *et al.*, 1999; Madhani *et al.*, 1999)]. None of these genes were required for viability of the *cdc24-4* mutant (*Fig. S1C*). To identify relevant polarity targets of the fMAPK pathway, GO term analysis was performed on data generated in (Chow *et al.*, 2019). Polarity targets of the fMAPK pathway included genes that regulate bud-site selection (*Fig. S1D*, *AXL2*, *BUD8, RSR1* and *RAX2*), polarity establishment (*MSB2*, *GIC2*, and *RGA1*), and septin ring organization (*GIC2, AXL2,* and *RGA1*). *GIC2* in particular encodes a direct effector of Cdc42p that along with *GIC1* functions in bud emergence [**Fig. 1A**, (Brown *et al.*, 1997; Chen *et al.*, 1997)]. The *GIC2* gene was a target of the fMAPK pathway [*Fig. S2A* and see (MacIsaac *et al.*, 2006)].

Gic2p was required for viability of the *cdc24-4* mutant [**Fig. 1B**, (Costanzo *et al.*, 2010)] and to suppress the growth defect of *cdc24-4* by p*STE11-4* (**Fig. 1B**). Expression of *GIC2* from a fMAPK pathway-independent promoter partly rescued the growth defect of the *cdc24-4* mutant (**Fig. 1B**, p*TEF2-GIC2*) even in cells lacking an intact fMAPK pathway (p*TEF2-GIC2 ste12*Δ). Gic2p played a unique role compared to Gic1p (*Fig. S2A* and *B*). Thus, Gic2p is a target of the fMAPK pathway required for fMAPK pathway-dependent rescue of the *cdc24-4* mutant.

Gic proteins regulate clustering or polarization of Cdc42p at bud sites (Daniels *et al.*, 2018; Kang *et al.*, 2018), which is a critical step in polarity establishment. Time-lapse fluorescence microscopy showed that GFP-Cdc42p clustered at a specific site on the cell cortex (**Fig. 1D**, wild type, asterisk), from which site new buds emerged. The *cdc24-4* mutant was defective for GFP-Cdc42p clustering and bud emergence (**Fig. 1D**, *cdc24-4*). Analysis of 30 cells showed this difference to be statistically significant (**Fig. 1E**). p*STE11-4* partially restored GFP-Cdc42p clustering and bud emergence to the *cdc24-4* mutant (**Fig. 1, D** and **E**). Rescue of bud emergence by p*STE11-4* was dependent on Gic2p (**Fig. 1, D** and **E**).

During bud emergence, Gic proteins also regulate formation of the septin ring (Iwase *et al.*, 2006; Bi and Park, 2012; Okada *et al.*, 2013; Sadian *et al.*, 2013; Kang *et al.*, 2018). In wild-type cells, the septin Cdc12p-GFP was localized in a ring prior to bud emergence (**Fig. 1F**, arrows) and to the mother-bud neck in budding cells (**Fig. 1F**, asterisk). In the *cdc24-4* mutant, Cdc12p-GFP showed a punctate pattern on the cell cortex (**Fig. 1F**, red arrows). Based on the analysis of over 150 cells, the localization defect was statistically significant (**Fig. 1F**). p*STE11-4* partially rescued the septin localization defect of the *cdc24-4* mutant (**Fig. 1, F** and **G**), which was dependent on Gic2p (**Fig. 1, F** and **G**). The fMAPK pathway did not suppress the growth defect of the *cdc12-6* mutant, which indicates that it functions upstream of the septins themselves (*Fig. S2C*). Gic2p also promotes the interaction between Cdc42p and Bni1p (Chen *et al.*, 2012). Bni1p was not required for GFP-Cdc42p clustering or bud emergence (**Fig. 1, E and F**) but was required for septin localization of the *cdc24-4* mutant (**Fig. 1, F** and **G**). Bni1p is required for septin ring assembly (Kadota *et al.*, 2004; Gladfelter *et al.*, 2005). In wild-type cells Gic2p was required for invasive growth (PWA, *Fig. S3A*), colony ruffling (*Fig. S3B*), and septin organization (*Fig. S3C-E*) of filamentous cells. Gic2p did not regulate the fMAPK pathway (*Fig. S3F*). These results establish a new role for the fMAPK pathway in regulating bud emergence by a mechanism that involves Gic2p.

### The fMAPK Pathway Regulates GTP-Cdc42p Levels During Filamentous Growth

Like other monomeric GTPases, the exchange of GDP for GTP alters the conformation of the protein and allows the protein to bind effector proteins. During budding, the increase in GTP-Cdc42p levels is critical for symmetry breaking and polarity establishment (Irazoqui *et al.*, 2003; Kozubowski *et al.*, 2008). Msb2p interacts with GTP-Cdc42p to activate the fMAPK pathway (Cullen *et al.*, 2004) and during filamentous growth, the level and activity of Msb2p is stimulated by positive feedback (*Fig. S1D*). To test whether Msb2p and the fMAPK pathway impacts the levels of GTP-Cdc42p in the cell, a fluorescent reporter that measures GTP-Cdc42p levels was examined [Gic2p-PBD-tdTomato (Okada *et al.*, 2013)]. Compared to cells undergoing yeast-form growth (YEPD), cells undergoing filamentous growth showed elevated levels of GTP-Cdc42p (**Fig. 2A**; YEP-GAL). The increase in GTP-Cdc42p levels was dependent on the fMAPK pathway (**Fig. 2A**) in a manner that was statistically significant (**Fig. 2B**, YEP-GAL; p-value < 0.00001). As previously shown (Okada *et al.*, 2013), the level of GTP-Cdc42p was also higher in cells responding to the mating pheromone α-factor (**Fig. 2, A** and **B**). In this case, the increase was independent of the fMAPK pathway.

**Figure 2.**
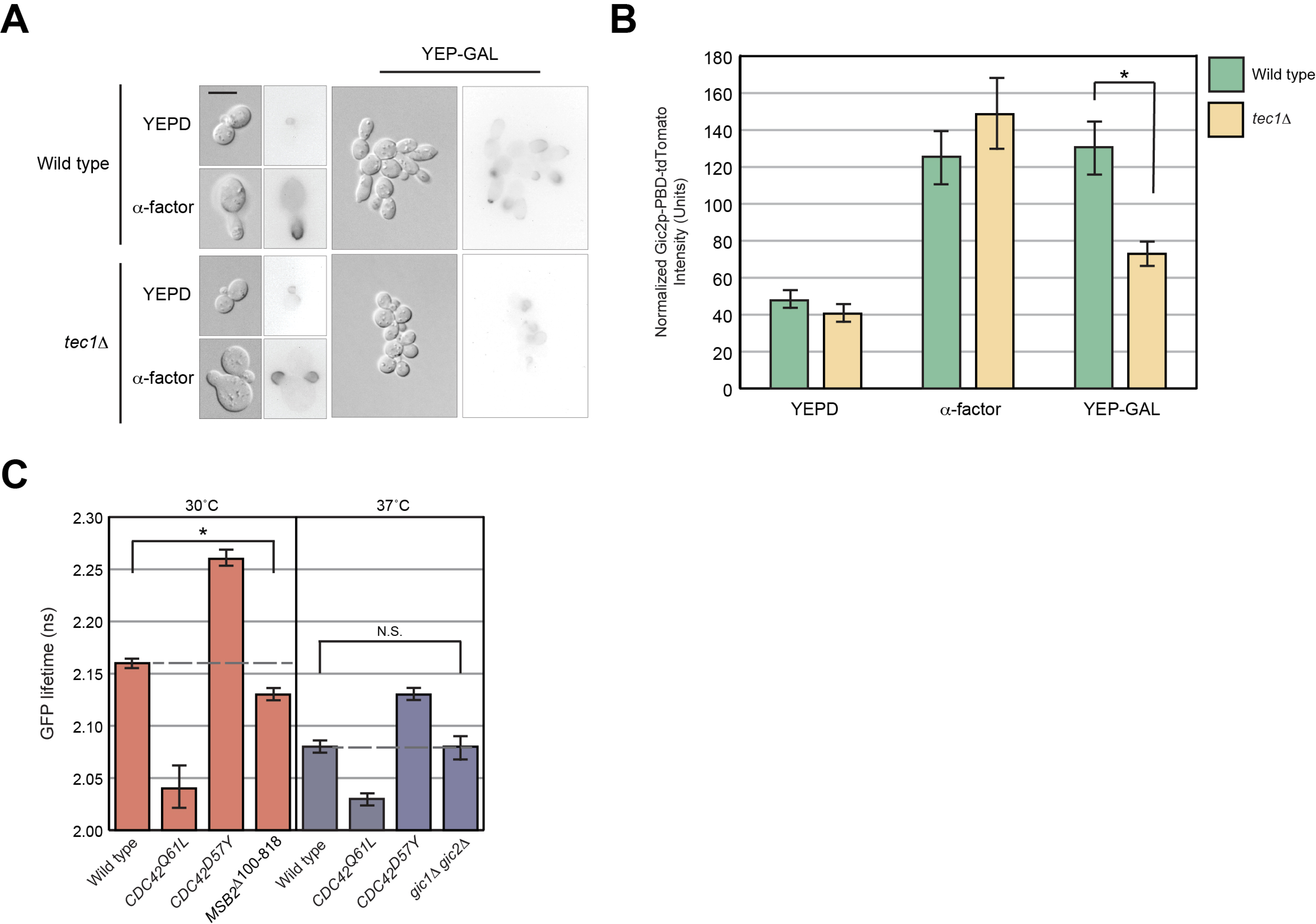
Role of the fMAPK pathway in regulating GTP-Cdc42p levels at site of bud emergence. (**A**) Localization of Gic2p-PBD-tdTomato in wild-type cells and the *tec1*Δ mutant under the indicated conditions. Micrographs were taken at the same exposure. Bar, 7.5 microns. (**B**) Quantitation of normalized total pixel intensity of Gic2p-PBD-tdTomato cluster in wild-type and *tec1*Δ mutant cells. Error bars represent standard error of mean of over 60 cells. Asterisk, p-value < 0.00001. (**C**) Wild-type cells expressing plasmid-borne Cdc42p biosensors and combinations of *MSB2*^Δ100-818^ examined by FLIM-FRET microscopy. In a separate experiment, wild type and the *gic1*Δ *gic2*Δ double mutant expressing the wild-type Cdc42p biosensor were grown to mid-log and shifted to 37°C for 4 h followed by examination by FLIM-FRET microscopy. Each histogram represents the mean of a set of measurements from three independent trials. The average of > 15 cells is reported for each strain. Error bars represent the standard error of mean. Asterisk, p-value <0.05. N.S., not significant.

One caveat in interpreting the above results is that *GIC2*-*PBD*-tdTomato is driven by its endogenous promoter, which, as shown above is regulated by the fMAPK pathway. As a separate test, the activity of a Cdc42p biosensor (Smith *et al.*, 2013) was examined by fluorescence lifetime imaging [FLIM-FRET (Sun *et al.*, 2011)]. In wild-type cells, the Cdc42p biosensor exhibited shorter lifetimes with a version that mimics the GTP-bound conformation (**Fig. 2C**, Cdc42p^Q61L^) and longer lifetimes with a version that mimics the GDP-bound conformation (**Fig. 2C**, Cdc42p^D57Y^). Activation of the fMAPK pathway reduced the lifetime of the biosensor (**Fig. 2C**, *MSB2*^Δ100-818^). This change corresponded to a 25% increase in the total levels of GTP-Cdc42p, which has previously been shown to reflect changes in Cdc42p activity during bud emergence (Smith *et al.*, 2013). Cdc42p activity can also be impacted by the Gic proteins (Kawasaki *et al.*, 2003; Daniels *et al.*, 2018; Kang *et al.*, 2018) but the lifetime of the biosensor was not altered in the *gic1*Δ *gic2*Δ double mutant (**Fig. 2C**). Hyperactive versions of Msb2p, Sho1p, and Ste11p also partially suppressed the growth defect of the *gic1*Δ *gic2*Δ double mutant at 37°C [*Fig. S4*, (Gandhi et al., 2006)] which supports a role for fMAPK in regulating bud emergence that is separate from Gic protein function.

### fMAPK Pathway Is Temporally and Spatially Regulated to Coincide with Bud Emergence

Bud emergence occurs in the G_1_ phase of the cell cycle (Hartwell *et al.*, 1970; Lew and Reed, 1993; Pringle *et al.*, 1995; Gulli *et al.*, 2000; Howell and Lew, 2012; Moran *et al.*, 2019). Whether the fMAPK pathway is active in the G_1_ phase of the cell cycle has not been explored. To address this question, cells were synchronized in the G_1_ phase of the cell cycle by α-factor (Breeden, 1997), and fMAPK pathway activity was assessed under basal (YEPD) and activating conditions (YEP-GAL) as cells progressed through the cell cycle. An epitope-tagged cyclin, Clb2p-HA, showed the expected pattern of cell-cycle regulation (Richardson *et al.*, 1992; Irniger *et al.*, 1995; Wasch and Cross, 2002; Cross *et al.*, 2005; Eluere *et al.*, 2007; Kuczera *et al.*, 2010; Cepeda-Garcia, 2017), increasing after α-factor release by 60 min and decreasing by 90 min as cells entered anaphase (**Fig. 3A**, α-HA). P∼Kss1p levels also increased after α-factor release by 80 min and peaked by 100 min (**Fig. 3A**; P∼Kss1p). By comparison, the level of P∼Fus3p, which is the MAP kinase that regulates the mating pathway, decreased after release from α-factor (**Fig. 3A**; P∼Fus3p). Conditions that activate the fMAPK pathway (YEP-GAL) led to a delay in Clb2p-HA accumulation (Leitao and Kellogg, 2017) and also showed an increase in P∼Kss1p levels in the G_2_/M phase of the cell cycle, prior to next round of bud emergence (**Fig. 3B**). These results demonstrate that the activity of the fMAPK pathway is cell-cycle regulated and increases prior to and during bud emergence.

**Figure 3.**
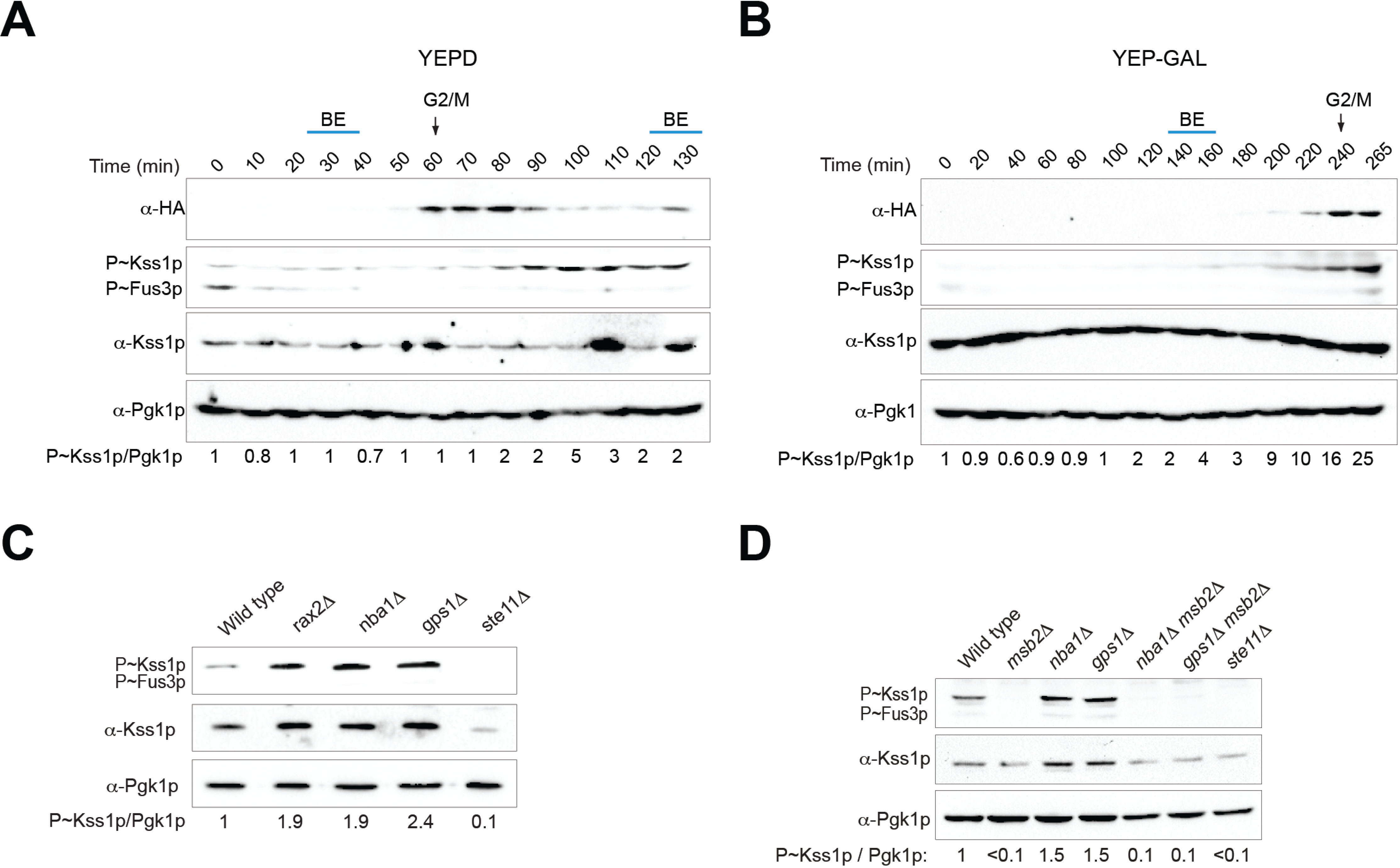
The activity of the fMAPK pathway is cell cycle regulated and inhibited at dormant sites. **(A)** Immunoblot analysis of wild-type cells synchronized in G_1_ by α-factor arrest and released in YEPD. Cell extracts were probed at indicated time points with antibodies to Clb2p-HA (α-HA), P∼Kss1p (α-p44/42), and Pgk1p, as a control for protein levels. Numbers refer to the ratio of P∼Kss1p to Pgk1p relative to 0 min, which was set to 1. BE, bud emergence. (**B**) Same as panel A, except cells were released into YEP-GAL media. (**C**) Immunoblot analysis of wild type and mutant combinations to assess fMAPK pathway activity. Pgk1p, loading control. Numbers refer to the ratio of P∼Kss1p to Pgk1p relative to wild type, set to 1. **(D)** Immunoblot analysis of the role of Msb2p in regulating the elevated fMAPK pathway activity in mutants lacking the negative polarity complex. See panel 3C for details.

Bud emergence is spatially regulated in that it results from Cdc42p activation at bud sites (Caviston *et al.*, 2003; Das *et al.*, 2007; Freisinger *et al.*, 2013; Okada *et al.*, 2013). We previously showed that bud-site-selection proteins regulate the fMAPK pathway (Basu *et al.*, 2016), which indicates that the fMAPK pathway is active at sites where bud emergence occurs. In line with this possibility, key regulators of the fMAPK pathway, including Sho1 (see below) and Ste20p are recruited to bud sites prior to bud initiation (Moran *et al.*, 2019). Moreover, polarity targets of the fMAPK pathway included proteins that promote budding at bud sites (*Fig. S1D*). Polarity targets also included proteins that prevent budding at previous division sites or cytokinesis remnants. These included Rga1p (*Fig. S1D*), which along with other GTPase activating proteins (GAPs) prevents budding within the existing growth site (Kadota *et al.*, 2004; Tong *et al.*, 2007; Miller *et al.*, 2017), and Rax2p (*Fig. S1D*), which restricts budding at previous division sites in a complex with Rax1p, Gps1p, Nba1p and Nis1p (Meitinger *et al.*, 2014). Rga1p has been shown to negatively regulate the fMAPK pathway (Smith *et al.*, 2002). Cells lacking Rax2p, Nba1p, or Gps1p also showed elevated fMAPK pathway activity (**Fig. 3C**, *Fig. S5A*), which indicates that the negative polarity complex negatively regulates the fMAPK pathway. The elevated activity of the fMAPK pathway in negative polarity complex mutants also required Msb2p (**Fig. 3D**, *Fig. S5A-B*). Cells lacking the negative polarity complex did not impact mating (*Fig. S5C*). Therefore, proteins that spatially promote budding promote fMAPK activity, and proteins that spatially restrict budding also restrict fMAPK pathway activity. Collectively, the data indicate that fMAPK pathway activity is temporally and spatially regulated to coincide with bud emergence.

### fMAPK Pathway Stimulates the Rate of Bud Emergence During Filamentous Growth

To further define how the fMAPK regulates bud emergence, the timing of bud emergence was examined using a septin marker (Cdc3p-mCherry) that shows a characteristic localization pattern throughout the cell cycle (Kim *et al.*, 1991; Lippincott *et al.*, 2001). In yeast-form cells grown in SD-URA (GLU), the timing of GFP-Cdc42p clustering, septin recruitment by Cdc3p-mCherry, and bud emergence were similar between wild-type cells and an fMAPK pathway mutant (*Fig. S6A*). In filamentous wild-type cells grown in S-GAL-URA (GAL), GFP-Cdc42p clustering occurred at the incipient site by 10 min [**Fig. 4; A** (green arrowheads) and **B**; **Movie 1**; *Fig. S7A*] from the preceding round of cytokinesis marked by the septin hourglass split into double ring [(**Fig**. **4A**, set as t=0) (Cid *et al.*, 2001; Lippincott *et al.*, 2001)]. The recruitment of Cdc3p-mCherry had also occurred within this duration [**Fig. 4; A** (red arrowheads) and **B**; **Movie 1**; *Fig. S7A*). The average time for bud emergence for 17 wild-type cells was recorded as 30 minutes from the preceding cytokinesis (**Fig. 4B**). Strikingly, in the fMAPK pathway mutant grown under filamentous conditions (GAL), delay in GFP-Cdc42p clustering, Cdc3p-mCherry recruitment and bud emergence were observed (**Fig. 4B**). For a given fMAPK mutant cell, initial GFP-Cdc42p clustering and Cdc3p-mCherry recruitment occurred within the same time span [(**Fig. 4, B** and **C** (example 1, green and red arrowheads)]. However, some cells showed disappearance and reappearance of the GFP-Cdc42p cluster at the initial polarity site [(**Fig. 4C** and **D** (oscillation seen in the pixel intensity graphs); **Movie 2-4**; *Fig. S7B-D*]. The disappearance and reappearance of the GFP-Cdc42p cluster, referred here as transient disappearance of polarity complex, accounted for a significant delay in bud emergence in the fMAPK pathway mutant (**Fig. 4, B** and **C**). Some of these cells even failed to make buds within the time of the experiment (**Fig. 4D**; **Movie 3-4**; *Fig. S7C-D*). In other examples, the initial polarity site disappeared and reappeared at a new site from where the bud came out [**Fig. 4E** (in the mother cell, compare the location of green and red arrowhead with the black arrow that marks bud emergence); **Movie 5-6**; *Fig. S7E-F*]. The transient disappearance of the polarity complex, seen at some level in wild-type cells (although they always made a bud), was increased by 3-fold in the fMAPK mutant (**Fig. 4F**). Representing GFP-Cdc42p intensity in each cell as the coefficient of variation (CV) of pixel intensity (Lai *et al.*, 2018) over time brought out the polarity defect in the fMAPK mutant relative to the wild type (**Fig. 4G**). The coefficient of variation (CV) of pixel intensity also showed a larger change in wild type (0.16 to 0.23 = 7) compared to the *ste12*Δ mutant (0.14 to 0.18 = 4), which was statistically significant (p-value, <0.01). These results define a function for the fMAPK pathway in stimulating the rate of bud emergence under conditions that promote filamentous growth.

**Figure 4.**
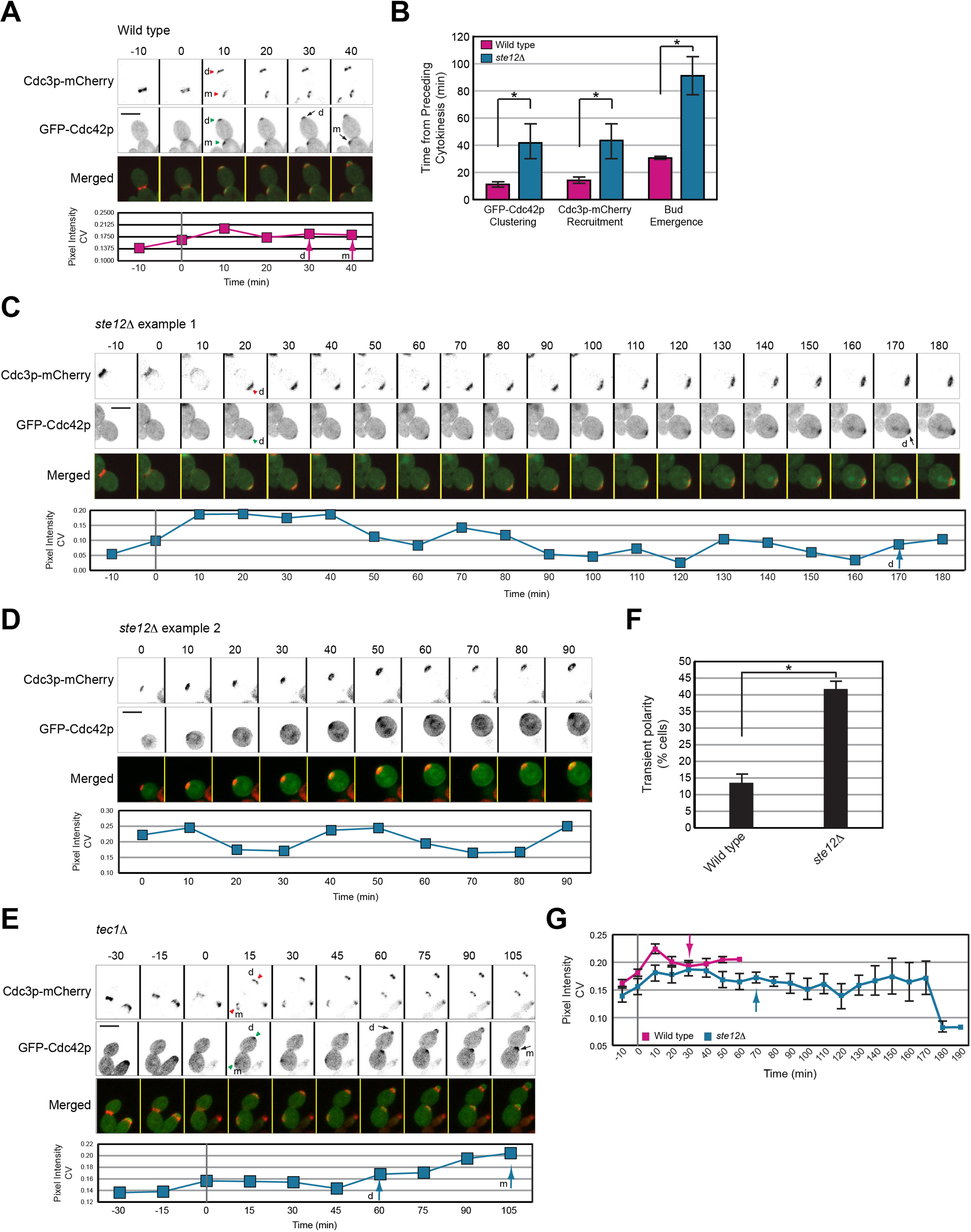
Role of the fMAPK pathway in regulating bud emergence during filamentous growth. (**A**) Inverted maximum intensity projection of GFP-Cdc42p clustering and bud emergence in wild-type cells in GAL. Cdc3p-mCherry was used as a marker for cell cycle progression. Septin hourglass split into double ring was set as time 0. Green arrowheads, first visible cluster of GFP-Cdc42p. Red arrowheads, recruitment of Cdc3p-mCherry at incipient site. Arrows, bud emergence. Graph represents GFP-Cdc42p intensity over time measured as coefficient of variance (CV) of pixel intensity of the entire cell. Arrow, timing of bud emergence. Bar, 5 microns. m, mother cell; d, daughter cell. (**B**) Quantitation of the timing of GFP-Cdc42p clustering, Cdc3p-mCherry recruitment and bud emergence in wild-type cells and *ste12*Δ cells grown on semi-solid S-GAL-URA media. Average of 17 cells for each strain is reported. Error bar represent standard error of mean. Asterisk, p-value <0.05. (**C**) Same as A, except *ste12*Δ mutant. (**D**) Same as C, except t=0 represents start of experiment. Time points for the preceding cell cycle were not available for this cell. (**E**) Same as A, except *tec1*Δ mutant. (**F**) Percent cells showing transient appearance and disappearance of polarity complex detected by co-localization of GFP-Cdc42p and Cdc3p-mCherry in cells from B. Error bar represents standard error of mean of 17 cells. Asterisk, p-value < 0.01. (**G**) GFP-Cdc42p intensity over time measured as pixel intensity CV for indicated strains. The average of 17 cells is reported for each strain. Error bars represent the standard error of mean.

To look at active Cdc42p during bud emergence in filamentous conditions, Gic2p-PBD-tdTomato reporter was co-localized with Sho1p-GFP, an fMAPK pathway regulator which is a direct effector of Msb2p (Cullen *et al.*, 2004; Tatebayashi *et al.*, 2007) that interacts with Cdc24p (Vadaie *et al.*, 2008), Ste20p, and Ste11p (Zarrinpar *et al.*, 2004; Tatebayashi *et al.*, 2006). Although Sho1p is known to localize to polarized sites (Pitoniak *et al.*, 2009), its localization throughout the cell cycle has not been explored. In wild-type cells, Sho1p-GFP was localized to incipient bud sites prior to bud emergence together with GTP-Cdc42p (*Fig. S6B-C*; wild type; red arrow). After bud emergence, Sho1p-GFP was found at the bud tips as has been reported (Pitoniak *et al.*, 2009). In *tec1*Δ mutant cells, Sho1p-GFP initially localized to the incipient bud site but failed to become enriched, and instead migrated back and forth along the distal pole (*Fig. S6C*; *tec1*Δ; black arrows). The low level of active Cdc42p in these cells rapidly disappeared. Bud formation was delayed and in the cell shown bud emergence did not occur within the time of the experiment. Thus, although, active Cdc42p clusters at incipient bud sites, an intact fMAPK pathway is required to promote bud emergence under nutrient limiting conditions. Cells lacking the fMAPK pathway showed a growth defect under filamentous conditions (*Fig. S6D*, GAL) and a defect in the rate of bud formation (*Fig. S6*, *E*, and *F*). In particular, at 5 h, 10 h, and 18 h, the *ste12*Δ mutant formed buds at a slower rate than wild-type cells. Therefore, the fMAPK pathway stimulates the rate of budding during filamentous growth.

### Activation of the fMAPK Pathway Induces Growth at Multiple Sites

Wild-type cells normally grow at a single site due to a regulatory phenomenon known as singularity in budding. Cells containing active versions of Cdc42p bypass this regulation and grow at multiple sites (Richman and Johnson, 2000; Caviston *et al.*, 2002; Wedlich-Soldner *et al.*, 2003; Wedlich-Soldner *et al.*, 2004; Howell *et al.*, 2009; Howell *et al.*, 2012). Cells containing hyperactive versions of Msb2p also had this property [**Fig. 5A**, (Basu *et al.*, 2016)]. Further inspection showed that 16% of cells carrying *MSB2*^Δ100-818^ showed multiple growth sites (**Fig. 5B**, *MSB2*^Δ100-818^). Given that the fMAPK pathway regulates bud emergence, the ability of Msb2p to induce multiple growth sites might be mediated by the fMAPK pathway. We found that the formation of multiple growth sites by *MSB2*^Δ100-818^ required the fMAPK pathway (**Fig. 5B**, *MSB2*^Δ100-818^ *ste12*Δ). Growth at multiple sites was also induced by *STE11-4* or overexpression of *SHO1* (**Fig. 5B**). The formation of multiple growth sites correlated with fMAPK pathway activity (**Fig. 5, B** and **C**). The activity of the fMAPK pathway is stimulated by positive feedback, which is evident by immunoblot of the Kss1p protein (**Fig. 5C**, middle blot), whose levels are controlled by the fMAPK pathway (Roberts *et al.*, 2000). Thus, Msb2p might itself induce multiple growth sites but require the fMAPK pathway for positive feedback. However, this was not the case. *MSB2*^Δ100-818^ expressed from an inducible promoter (*pGAL*) that did not depend on the fMAPK pathway also required Ste12p to form multiple growth sites (**Fig. 5, B** and **C**, compare *GAL-MSB2*^Δ100-818^ to *GAL-MSB2*^Δ100-818^ *ste12*Δ). Therefore, Msb2p activation itself is not sufficient to induce growth at multiple sites. The fact that *MSB2*^Δ100-818^ is more effective at inducing growth at multiple sites than *STE11-4* can be explained by fMAPK pathway activity.

**Figure 5.**
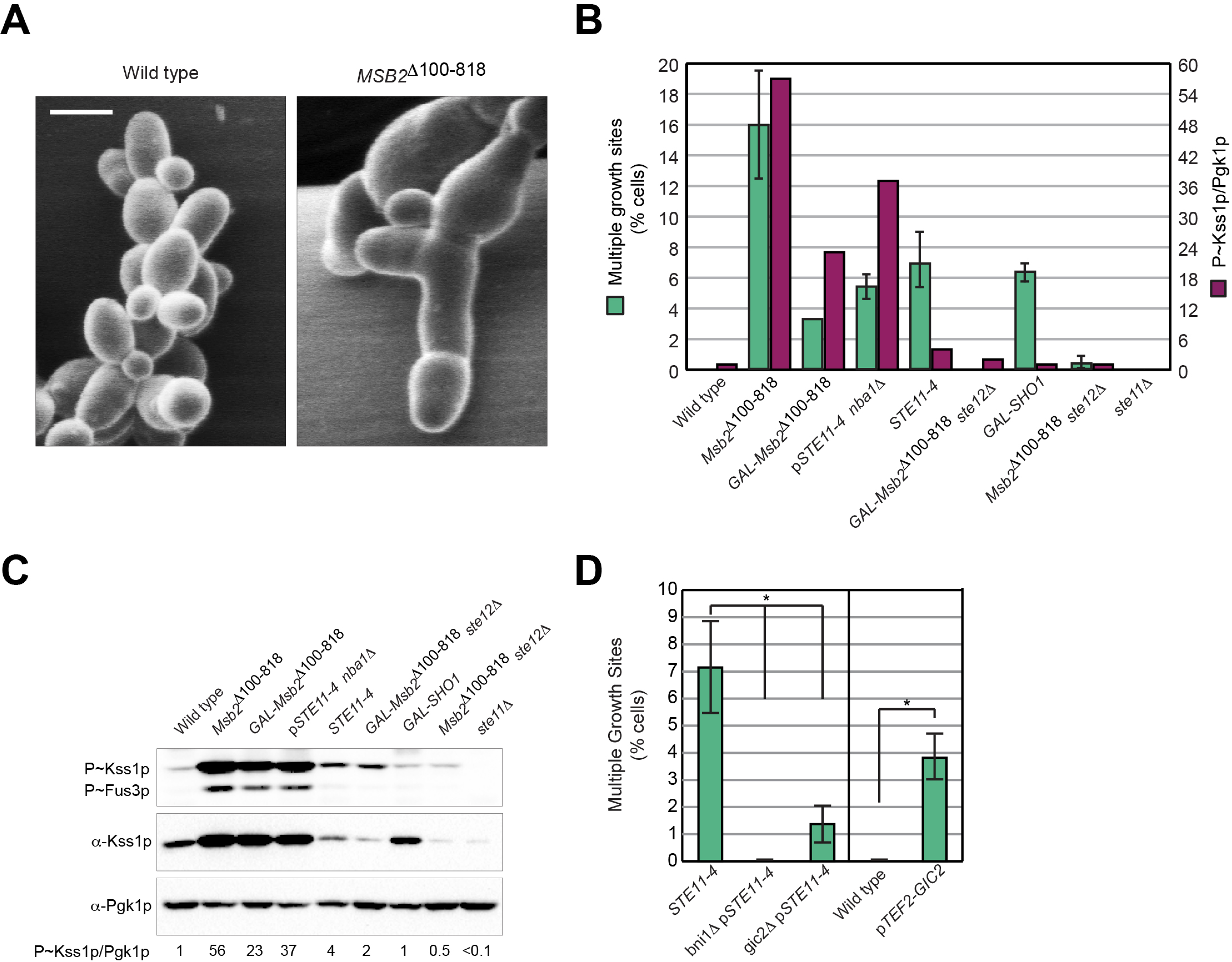
Role of the fMAPK pathway in making multiple growth sites. (**A**) S.E.M images of wild type and *MSB2*^Δ100-818^ examined for multiple growth sites. Bar, 5 microns. (**B**) Comparison of percentage of multiple growth sites by the single cell assay to P∼Kss1p levels in panel C in the indicated strains. Error bars represent standard error of mean from three separate trials. Over 50 cells were counted in each trial. The histogram for P∼Kss1p represents the ratio of P∼Kss1p to Pgk1p relative to wild type, which was set to 1. (**C**) Immunoblot analysis of fMAPK activity in the indicated strains grown in YEP-GAL. Pgk1p, loading control. Numbers refer to the ratio of P∼Kss1p to Pgk1p relative to wild type, which was set to 1. (**D**) Percentage of multiple buds by the single cell assay in indicated strains. In a separate experiment, wild-type cells expressing *TEF2-GIC2* were grown to saturation and evaluated for the percentage of multiple buds. See panel B for details. Asterisk, p-value < 0.01.

Transcriptional targets of the fMAPK pathway may be required for the formation of multiple growth sites. Bni1p and Gic2p were required for multiple growth site formation by the fMAPK pathway (**Fig. 5D**). High-copy expression of *GIC2* itself induced multiple growth sites (**Fig. 5D**), in line with a previous report (Jaquenoud *et al.*, 1998). The negative polarity complex did not impact growth at multiple sites (**Fig. 5B**, p*STE11-4 nba1*Δ), although it did stimulate the fMAPK pathway activity (**Fig. 5C**). Collectively, these results support a role for the fMAPK pathway in the regulation of bud emergence, because activation of the fMAPK pathway can induce growth at multiple sites.

### The fMAPK Pathway Induces Symmetry Breaking in the Growing Bud

The multiple growth sites produced by the fMAPK pathway differed in shape from previous reports on multiple bud formation by hyper-activation of Cdc42p (Caviston *et al.*, 2002; Wedlich-Soldner *et al.*, 2003; Wedlich-Soldner *et al.*, 2004). Hyperactive Cdc42p induces symmetry breaking in the mother cell to trigger formation of a second bud. By comparison, the hyperactive fMAPK pathway did not impact budding in mother cells. The multiple growth sites were formed inside the growing bud (**Fig. 6A**). To our knowledge, this new phenotype has not been previously characterized. Growth at a second site within the growing bud was not due to abortive growth at the initial site, because polarity proteins including Cdc24p-GFP (**Fig. 6B**) and F-actin (**Fig. 6C**, *Fig. S8A-B*) localized to multiple sites. Cells expressing *MSB2*^Δ100-818^ showed asymmetric clustering of GFP-Cdc42p (**Fig. 6D**, **Movie 8**, *Fig. S9B*). Over time, GFP-Cdc42p migrated along the bud cortex (**Fig. 6D**), which was evident by kymograph analysis (**Fig. 6E**), which allows tracking of changes in protein localization over time (Kaksonen *et al.*, 2003). Sec3p-GFP localization corroborated that these sites were actively growing (*Fig. S8B*). Some wandering of GFP-Cdc42p occurred in wild-type cells (**Movie 7**, *Fig. S9A*), which might be due to the off-center delivery of vesicles that dilute the polarity complex (Dyer *et al.*, 2013; Chiou *et al.*, 2017).

**Figure 6.**
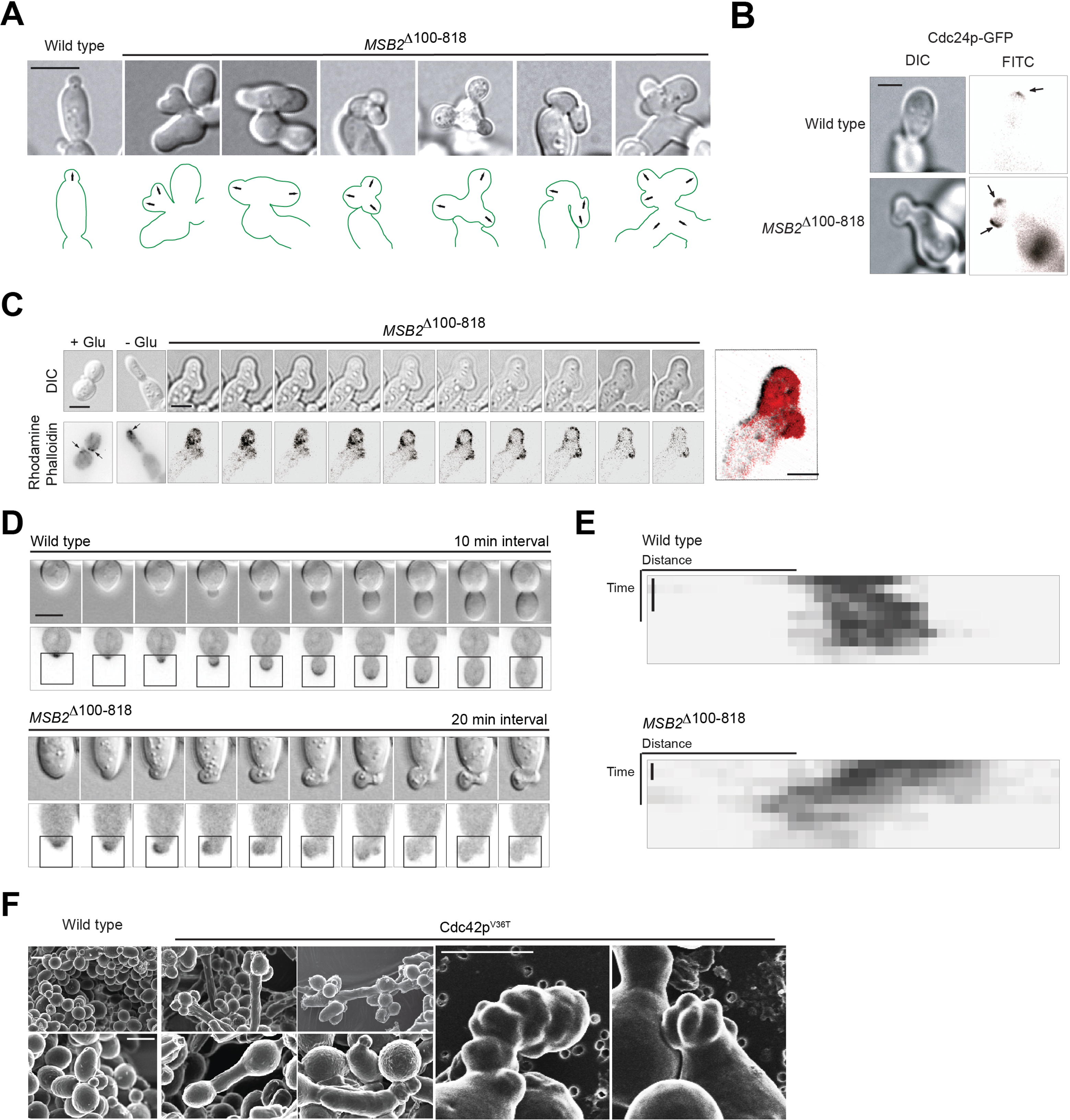
Activation of the fMAPK pathway leads to wandering of polarity. (**A**) Wild-type cells and cells carrying *MSB2*^Δ100-818^ were examined on S-GLU media by the single cell invasive growth assay. Arrows indicate growth sites. Bar, 5 microns. (**B**) Wild type and *MSB2*^Δ100-818^ containing Cdc24p-GFP were examined by single cell invasive growth assay. Arrows points to Cdc24p-GFP at the growing tip. Scale bar, 5 microns. (**C**) At left, actin staining of cells by rhodamine phalloidin. Bar, 5 microns. Micrographs for *MSB2*^Δ100-818^ shown at different focal planes in the Z-axis. At right, inverted maximum intensity projection of cell on left shown using red lookup table. (**D**) Time series of wild type and *MSB2*^Δ100-818^ expressing GFP-Cdc42p evaluated for multiple growth sites. Bar, 5 microns. Time interval; WT, 10 min; *MSB2*^Δ100-818^, 20 min. Highlighted regions were used to prepare kymographs in D. DIC and inverted maximum intensity projection shown. **(E)** Kymographs were prepared using ImageJ as described in materials and methods. Bar, 40 min. (**F**) SEM images of wild type and Cdc42p^V36T^. Scale bar, 5 microns.

The hyperactive fMAPK pathway may alter the dynamics of the interaction between Cdc42p and its effectors in a manner that makes the Cdc42p polarity axis ‘forget’ its orientation. This may mimic symmetry breaking occurring but within the bud cortex. In line with this possibility, versions of Cdc42p that contain point mutations in the effector-binding domain (such as Cdc42p^V36T^), which impair its interaction with effector proteins (Gladfelter *et al.*, 2001; Gladfelter *et al.*, 2002; Gladfelter *et al.*, 2005), showed a similar phenotype (**Fig. 6F**). Here multiple protrusions occurred adjacent to the previous polarization site. Similarly, cells expressing *MSB2*^Δ100-818^ formed multiple growth projections (**Fig. 6A**, three, four, or five sites). Other genes can induce multiple buds when overexpressed (Sopko *et al.*, 2006), but these did not rescue the growth defect of the *cdc24-4* mutant and in most cases worsened it (*Fig. S10*). These results demonstrate the importance of a properly regulated fMAPK pathway in bud morphogenesis.

## DISCUSSION

Here we show that bud emergence, which is one the most intensively studied and well understood polarity establishment processes in eukaryotes, is regulated by a MAPK pathway. The fMAPK pathway regulates transcription of polarity target genes and GTP-Cdc42p levels to increase the rate of bud emergence during filamentous growth. In this way, bud emergence is regulated by a pathway whose activity is sensitive to extrinsic cues. Importantly, the ability of the fMAPK pathway to induce the expression of polarity target genes allows the pathway to tailor bud emergence by altering the levels of proteins that act at multiple steps in the polarity pathway.

The fMAPK pathway may also regulate bud emergence during vegetative growth. Ste20p is the first effector recruited by Cdc42p at bud sites (Moran *et al.*, 2019), which is known to activate the fMAPK pathway at bud sites to regulate bud emergence. Ste12p and Tec1p are also required for viability of the *cdc24-4* mutant under vegetative growth conditions. A function for the fMAPK pathway in regulating bud emergence might be masked by genetic buffering under normal growth conditions. Indeed, cells lacking *TEC1* are synthetically lethal with a diverse class of cytoskeletal and cell-cycle regulatory genes (Costanzo *et al.*, 2010). More recently, Rsr1p in its GDP-locked state has been shown to regulates the timing of Cdc42p polarization in the early G_1_ by interaction with Bem1p (Miller *et al.*, 2019). Rsr1p (Basu *et al.*, 2016) and Bem1p (Basu *et al.*, UNDER REVIEW) both regulate the fMAPK pathway.

A key polarity target of the fMAPK pathway is Gic2p, which is a direct effector of Cdc42p that controls multiple steps in bud emergence. The regulation of *GIC2* expression may be a key step in the regulation of polarity establishment in general. *GIC2* expression is regulated by the fMAPK pathway and other proteins that regulate filamentous growth, including Phd1p (Gimeno and Fink, 1994; MacIsaac *et al.*, 2006), Rim101p (Lamb and Mitchell, 2003; Hu *et al.*, 2007), SAGA (Venters *et al.*, 2011) and Rpd3p(L) (Hu *et al.*, 2007; Venters *et al.*, 2011). *GIC2* expression is also regulated during mating (Roberts *et al.*, 2000) and the Gic proteins are required for shmoo formation (Brown *et al.*, 1997). Gic2p may function in other contexts as well, and has been implicated as an effector of the Protein Kinase C pathway (Zanelli and Valentini, 2005). Generally speaking, changes in gene expression may impact the regulation of bud emergence during cell differentiation and the response to stress.

Here and in Basu *et al*. (2016) we identify cross talk between the polarity pathway and the fMAPK pathway that impacts bud emergence and filamentous growth (**Fig. 7**). One new connection is that transcriptional amplification of fMAPK pathway target genes by positive feedback stimulates GTP-Cdc42p levels. A second connection is that bud-site-selection proteins that regulate the fMAPK pathway may reinforce bud emergence at bud sites, potentially by altering the expression of polarity target genes. A third connection is that the fMAPK pathway induces target genes that prevent growth and MAPK signaling at dormant growth sites (Meitinger *et al.*, 2013). Such cross talk reveals the precision that is required for proper bud emergence. Too little fMAPK pathway activity causes a delay in bud emergence during filamentous growth, and too much leads to growth at multiple sites. Precise regulation of the fMAPK pathway comes from activation at the right place (incipient bud sites) and time (at M/G_1_). The fact that the fMAPK pathway is cell-cycle regulated is a novel finding. Cell-cycle regulation of the fMAPK pathway might occur through *TEC1*, which like other G_1_ specific genes in the ‘SIC’ cluster (Cho *et al.*, 1998; Spellman *et al.*, 1998; Wittenberg and Reed, 2005) is induced at the M/G_1_ boundary by the cell-cycle regulated transcription factor Swi5p (Spellman *et al.*, 1998).

**Figure 7.**
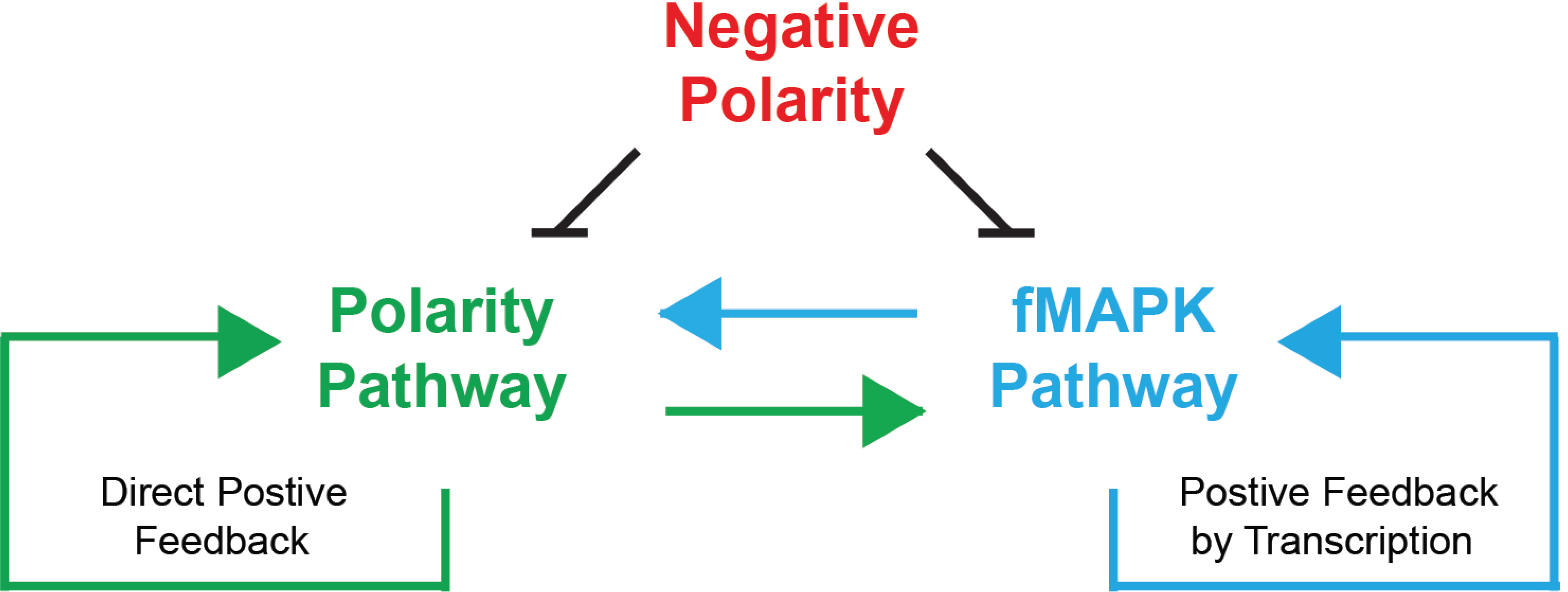
Model for the regulation of bud emergence that involves cross-regulation between the polarity and the fMAPK pathways. Arrows, positive regulation; bar, negative regulation.

During vegetative growth in *S. cerevisiae*, Cdc42p-dependent budding is dictated by bud-site-selection proteins (Chiou et al., 2017). In other fungal species, diverse mechanisms of growth control predominate. In *Schizosaccharomyces pombe*, growth at the two poles is maintained by regulated oscillations in Cdc42p activity (Das *et al.*, 2012). Changes in the orientation of the polarity complex occurs during mating (Nern and Arkowitz, 2000; Dyer *et al.*, 2013) and filamentous growth (this study) and may be a common feature of fungal cell differentiation. In the filamentous fungus *Ashbya gossypii*, Rsr1p regulates symmetric growth cone formation at the hyphal tip (Bauer *et al.*, 2004). In the major human fungal pathogen *Candida albicans*, Rsr1p and Cdc42p are also required for proper hyphal growth (Si *et al.*, 2016). MAPK pathways may be critical regulators of Cdc42p-dependent morphogenesis during hyphal/pseudohyphal growth. Parallels can also be drawn to higher eukaryotes. In mammals, the ERK pathway is directly involved in breaking radial symmetry of spreading RAT2 fibroblast cells (Klimova *et al.*, 2016). In this case, ERK spatially restricts p190A-RhoGAP activity to limit growth at the cell rear. Therefore, functional crosstalk between MAPK pathways and Rho GTPases may constitute an underexplored regulatory circuit in many systems.

## MATERIALS AND METHODS

### Strains and Plasmids

Yeast strains are described in Table 1. Plasmids are listed in Table 2. Gene disruptions and *GAL1* promoter fusions were made by PCR-based methods (Baudin *et al.*, 1993; Longtine *et al.*, 1998). Some gene disruptions were made with antibiotic resistance markers *KanMX6* (Longtine *et al.*, 1998), *HYG* and *NAT* (Goldstein and McCusker, 1999). Internal epitope fusions were made by the pop-in pop-out strategy (Schneider *et al.*, 1995). Some strains were made *ura3-* by selection on 5-fluoroorotic acid (5-FOA). Gene disruptions were confirmed by polymerase chain reaction (PCR) Southern analysis and confirmed by phenotype when applicable.

**Table 1.**
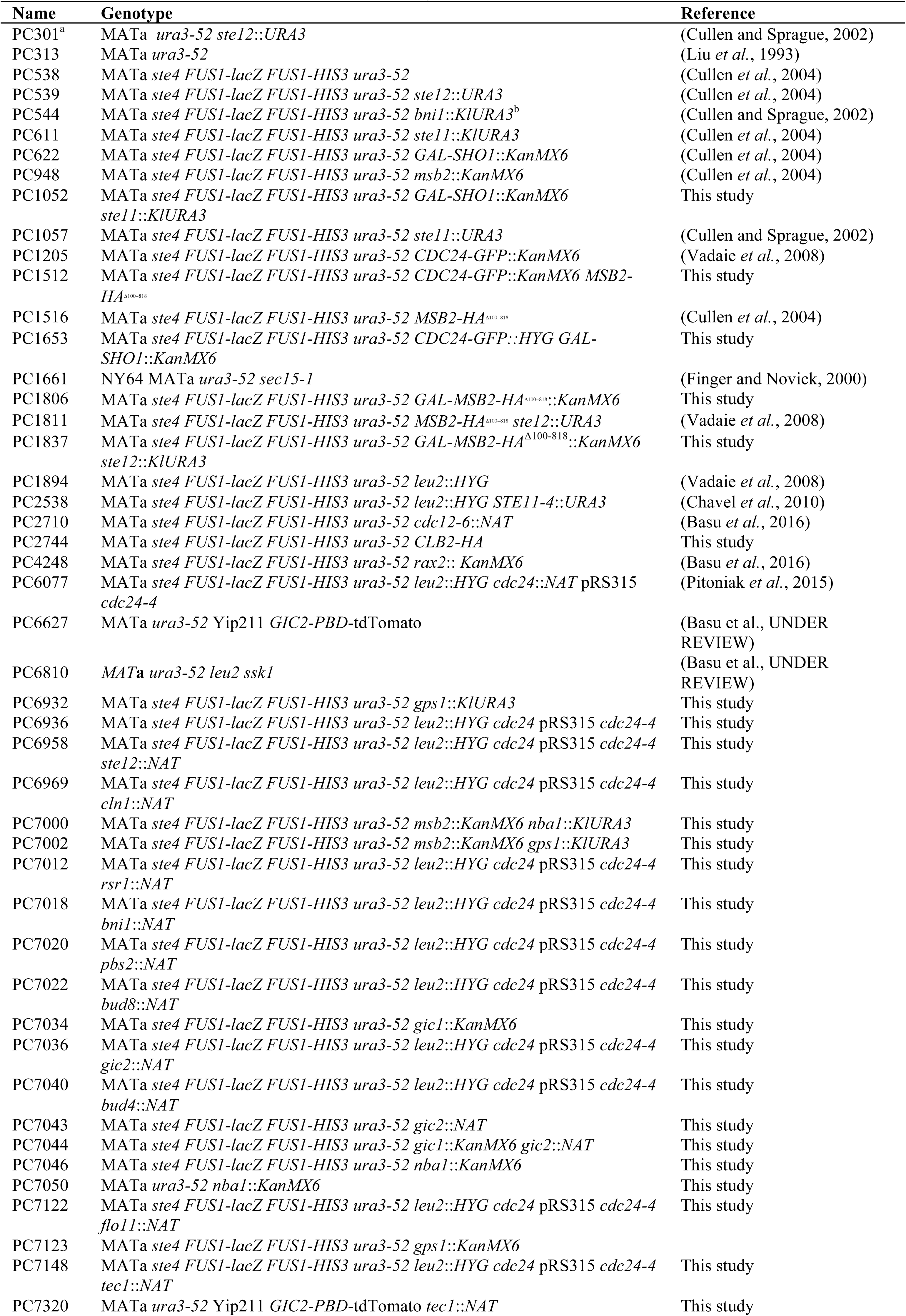
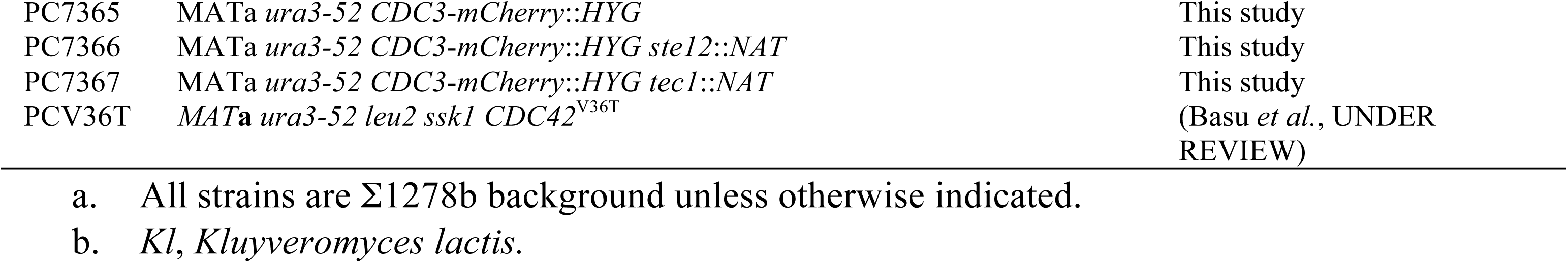
Yeast strains used in the study.

**Table 2.** Plasmids used in the study. ^a^.

| Name | Description | Reference |
| --- | --- | --- |
| PC187 | YCp50-*STE4* CEN/URA3 | (Stevenson *et al.*, 1992) |
| PC1333 | YCp50-*STE11-4* | (Stevenson *et al.*, 1992) |
| PC1365 | YCplac181 *CDC12-GFP* | (Fares *et al.*, 1996) |
| PC1665 | p*SEC3-GFP* | (Finger and Novick, 1997) |
| PC1696 | p*GFP-MSB2* | (Vadaie *et al.*, 2008) |
| PC1715 | pRS316-*SHO1*<sup>D16H P120L</sup>-*GFP* | (Vadaie *et al.*, 2008) |
| PC1885 | pMPY-3xHA | (Schneider *et al.*, 1995) |
| PC1964 | pRS316-*SHO1*<sup>D16H</sup>-*GFP* | (Marles *et al.*, 2004) |
| PC2151 | pRS315 | (Sikorski and Hieter, 1989) |
| PC2205 | pNAT | (Goldstein and McCusker, 1999) |
| PC2206 | pHYG | (Goldstein and McCusker, 1999) |
| PC2207 | pRS316 | (Sikorski and Hieter, 1989) |
| PC6365 | YEp352-*TEF2* | (Pitoniak *et al.*, 2015) |
| PC6454 | pRS316-*GFP-linker-CDC42* | (Basu *et al.*, UNDER REVIEW) |
| PC6626 | Yip211 *GIC2-PBD*-tdTomato | (Okada *et al.*, 2013) |
| PC7082 | YEp352-*TEF2-GIC2* | This study |
| PC7137 | pRS316- p*ACT1-EGFP-CRIBCLA4-CDC42*<sup>ΔCAAX</sup>-*mCherry-CAAX* | (Smith *et al.*, 2013) |
| PC7138 | pRS316- p*ACT1-EGFP-CRIBCLA4-CDC42*<sup>Q61LΔCAAX</sup>-*mCherry-CAAX* | (Smith *et al.*, 2013) |
| PC7139 | pRS316- p*ACT1-EGFP-CRIBCLA4-CDC42*<sup>D57YΔCAAX</sup>-*mCherry-CAAX* | (Smith *et al.*, 2013) |
| PC7143 | YCplac181 *CDC12-GFP::KanMX6* | This study |
| PC7150 | YCplac181 *CDC12-GFP::NAT* | This study |
| PC7340 | pRS316-*GFP-linker-CDC42::KanMX6* | This study |
| PC7357 | pRS316-*GFP-linker-CDC42::NAT* | This study |
| PC7359 | pRS316-p*ACT1-EGFP-CRIBCLA4-CDC42*<sup>ΔCAAX</sup>-*mCherry-CAAX::KanMX6* | This study |
| PC7361 | pRS316-*SHO1*<sup>D16H</sup>-*GFP::KanMX6* | This study |
| PC7363 | pRS316-*SHO1*<sup>D16H</sup>-*GFP::NAT* | This study |
| PC7381 | pRS315 *cdc24-4* | (Pitoniak *et al.*, 2015) |

a. The following plasmids were used from the movable ORF collection (Gelperin *et al.*, 2005): p*GAL-* *KIN3-HA*, p*GAL-LRE1-HA*, p*GAL-BET1-HA*, p*GAL-CLB4-HA*, p*GAL-YAP1801-HA*, p*GAL-YDL186W-* *HA*, p*GAL-DMA1-HA*, p*GAL-IQG1-HA*, and p*GAL-STE7-HA*.

The pRS series of plasmids (pRS315 and pRS316) have been described (Sikorski and Hieter, 1989). To construct *cdc24-4* mutant (PC6936), NAT cassette in PC6077 was replaced with *URA3* using pop-in pop-out strategy (Schneider *et al.*, 1995). Following complimentary primers were used-5’ CAGAAGAGTACCATTGCTGTTATCATTTGTTGCCTAGCCCTATCAACTAAAGGGAACAAAAGCTGG3’ and 5’ CAATATAATGGGTATAGCTTGAACATCTGCCCCTCTCTATCTAATGTCACTATAGGGCGAATTGG3’. Strain was subsequently made *ura3*Δ by selection on 5-fluoroorotic acid (5-FOA). *cdc24*Δ was confirmed by phenotypic analysis. Msb2p [p*GFP-MSB2* (Adhikari *et al.*, 2015)], Sho1p [pRS316-*SHO1^D16H,P120L^-GFP,* or *SHO1^P120L^* (Vadaie *et al.*, 2008)], and Ste11p [YCp50-*STE11-4* (Stevenson *et al.*, 1992)], have been described. Plasmid pRS316-*SHO1-GFP* has been described (Marles *et al.*, 2004) and was provided by Dr. Alan Davidson (University of Toronto, Ontario CA). In *SHO1*^D16H^, the D16H mutation alone does not hyperactivate the fMAPK pathway, but together with P120L mutation results in a hyperactive version of the protein. Overexpression constructs came from an ordered collection (Gelperin *et al.*, 2005). YEp352-*TEF2-GIC2* was created by amplifying the *GIC2* open reading frame (ORF) by PCR from genomic DNA using complimentary primers pairs 5’GACTCTAGAATGACTAGTGCAAGTATTACCAATACTGGAAACG3’ and 5’CAGTGTCGACTTAAGTTTGCAGGGGCTCGAGC3’. The PCR product was digested with XbaI and SalI and inserted into the YEp352-*TEF2* plasmid (Pitoniak *et al.*, 2015) digested with the same enzymes. Positive clones were confirmed by DNA sequencing.

Plasmid YCplac181-*CDC12-GFP::KanMX6* was constructed from YCplac181-*CDC12-GFP* (PC1365), provided by J. Pringle (Fares *et al.*, 1996), by homologous recombination. The *KanMX6* cassette was amplified by PCR with flanking sequences to *LEU2* locus on the YCplac181-*CDC12-GFP* plasmid. Complimentary primers were designed 5’-ATGTCTGCCCCTAAGAAGATCGTCGTTTTGCCAGGTGACCACGTTGCGGATCCCCGGGTTAATTA-3’ and 5’-TTAAGCAAGGATTTTCTTAACTTCTTCGGCGACAGCATCACCGACTGAATTCGAGCTCGTTTAAAC-3’ (Millipore Sigma, St. Louis, MO). The PCR product was co-transformed with YCplac181-*CDC12-GFP* digested with AflII to target integration at the *LEU2* locus. Plasmids were rescued from isolates that were *KanMX6* positive. YCplac181-*CDC12-GFP::NAT* was made by the same strategy.

pRS316-*GFP-CDC42::KanMX6* was made from pRS316-*GFP-CDC42* (PC6454) by targeting integration at the *URA3* locus using complimentary primers 5’-ATGTCGAAAGCTACATATAAGGAACGTGCTGCTACTCATCCTAGTCGGATCC CCGGGTTAATTA-3’ and 5’-TTAGTTTTGCTGGCCGCATCTTCTCAAATATGCTTCCCAGCCTGGAATTCGAGCTCGTTTAAAC-3’ (Millipore Sigma, St. Louis, MO). pRS316-p*ACT1*-*EGFP*-CRIB*CLA4*-*CDC42*^ΔCAAX^-mCherry-CAAX*::KanMX6* was made from pRS316-p*ACT1-EGFP*-CRIB*CLA4*-*CDC42*^ΔCAAX^-mCherry-CAAX (PC7137) (Smith *et al.*, 2013) and p*SHO1*^D16H^-*GFP::KanMX6* was made from p*SHO1*^D16H^-*GFP* (PC1964) by targeting integration at the *URA3* locus using same complimentary primers as for pRS316-*GFP-CDC42::KanMX6*. The PCR product was co-transformed with plasmids digested with BsmI to target integration at the *URA3* locus. Plasmids were rescued from isolates that were *KanMX6* positive. pRS316-*GFP-CDC42::NAT* and p*SHO1*^D16H^-*GFP::NAT* were made by the same strategy.

Plasmids were recovered from cells with Promega Wizard Plus SV Miniprep DNA Purification system (A1460, Promega, Madison, WI) used as specified by the manufacturer with following modification. 100 µl of glass beads were added at the lysis step, and cells were vortexed for 5 min before the addition of neutralization buffer. Rescued plasmids were transformed into *E. coli*, and positive clones were confirmed by DNA sequencing, by GENEWIZ (South Plainfield, NJ) and by fluorescence microscopy after transformation in yeast.

### Filamentous Growth and Mating Assays

Yeast and bacterial strains were manipulated by standard methods (Sambrook, 1989; Rose, 1990). The single cell invasive growth assay (Cullen and Sprague, 2000) and the plate-washing assay (Roberts and Fink, 1994) were performed as described. Actin staining by rhodamine phalloidin was performed as described (Yuzyuk and Amberg, 2003). In cells lacking an intact mating pathway (*ste4*Δ), the *FUS1-HIS3* reporter (McCaffrey *et al.*, 1987) was used to evaluate fMAPK pathway activity (Cullen *et al.*, 2004). *FUS1-HIS3* activity was measured by spotting equal concentrations of cells onto SD-HIS medium and SD-HIS medium containing 3-Amino-1,2,4-triazole (3-ATA).

Halo assays were performed as described (Jenness *et al.*, 1987). A saturated culture of cells (A_600_ = 0.1) was spread on YEPD media and allowed to dry. 3 µl and 10 µl α-factor (1mg/ml) were spotted on the plate. Plates were incubated at 30°C and photographed at 24 h and 48 h. Experiments were performed in triplicate. Halo diameter (in cm) was measured by ImageJ analysis and plotted as a function of α-factor concentration. For Lat-A sensitivity, saturated culture of cells (A_600_ = 0.1) was spread onto SD-URA-LEU media containing 0.5 M sorbitol. 10 µl of Lat-A (Sigma, L5163) at 0 (10 mM DMSO) 0.1 mM, 0.2 mM and 0.5 mM (in 10 mM DMSO) were spotted on plates. Halo diameter (in cm) was measured by ImageJ analysis and plotted as a function of Lat-A concentration.

### Evaluating MAP Kinase Phosphorylation by Immunoblot Analysis

Immunoblot analysis was used to detect phosphorylated MAP kinases as described (Sabbagh *et al.*, 2001; Lee and Dohlman, 2008; Basu *et al.*, 2016). In *cdc24-4* mutant combinations, cells were grown to mid-log in 10 ml SD-URA at 30°C and then incubated at 37°C for 4 h. Cells were harvested by centrifugation. Pellets were washed once with water and flash frozen in liquid nitrogen. In experiments that did not involve *cdc24-4*, cells were grown in YEPD or YEP-GAL media for the times incubated. Proteins were precipitated by trichloroacetic acid (TCA) and analyzed by sodium dodecyl sulfate polyacrylamide gel electrophoresis (SDS-PAGE) (10% acrylamide). Proteins were transferred to nitrocellulose membranes (Amersham^TM^ Protran^TM^ Premium 0.45 µm NC, GE Healthcare Life sciences, 10600003).

### Cell Synchronization and Cell Cycle Analysis of MAP Kinase Activity

Cell synchronization experiments were performed as described (Breeden, 1997). The strain harboring Clb2p-HA (PC2744) was transformed with a plasmid containing the *STE4* gene [p*STE4* (Stevenson *et al.*, 1992)]. Cells were grown to an optical density (O.D.) A_600_ of 0.2 in SD-URA media. Cells were washed and resuspended in equal volume of YEPD and incubated for 90 min at 30°C. α-factor was added to a final concentration of 5 µg/ml and the culture was incubated for 90 min to arrest cells in the G_1_ phase of the cell cycle. Arrested cells were washed twice with water and resuspended in fresh YEPD or YEP-GAL media to release cells into the cell cycle. 10 ml aliquots were harvested every 10 min, flash frozen in liquid nitrogen and stored at −80°C.

### Immunoblot Analysis

ERK-type MAP kinases (P∼Kss1p and P∼Fus3p) were detected using p44/42 antibodies (Cell Signaling Technology, Danvers, MA, 4370) at a 1:5,000 dilution. Kss1p was detected using α-Kss1p antibodies (Santa Cruz Biotechnology, Santa Cruz, CA; #6775) at a 1:5,000 dilution. Clb2p-HA was detected using the α-HA antibody at a 1:5,000 dilution (Roche Diagnostics, 12CA5). For secondary antibodies, goat anti-rabbit IgG-HRP antibodies were used at a 1:10,000 dilution (Jackson ImmunoResearch Laboratories, Inc., West Grove, PA, 111-035-144). Mouse α-Pgk1p antibodies were used at a 1:5,000 dilution as a control for total protein levels (Novex, 459250). Goat α-mouse secondary antibodies were used at a 1:5,000 dilution to detect primary antibodies (Bio-Rad Laboratories, Hercules, CA, 170-6516). Blots were visualized by chemiluminescence using a Bio-Rad ChemiDoc XRS+ system (Bio-Rad, 1708265). Quantitation of band intensities for immunoblot analysis was performed with Image Lab Software (Bio-Rad, Inc.). For blots to evaluate phosphorylated MAP kinase proteins, membranes were incubated in 1X TBST (10 mM TRIS-HCl pH 8, 150 mM NaCl, 0.05% Tween 20) with 5% BSA. For other immunoblots, membranes were incubated in 1X TBST with 5% non-fat dried milk. All primary incubations were carried out at 16 h at 4°C. Secondary incubations were carried out at 25°C for 1 h.

### Growth Assays for Temperature-Sensitive Mutants

Wild type and temperature-sensitive mutants containing desired plasmids were grown to saturation in SD-URA media. For each strain, O.1 O.D._600_ of cells were serially diluted 4 times in distilled water and spotted onto SD-URA-LEU+ 0.5M Sorbitol plates. The plates were incubated at 30°C and 37°C. For *cdc12-6* suppression, cells were spotted onto SD-URA plates and incubated at 24°C and 30°C since the *cdc12-6* mutant had a severe growth defect at 30°C and failed to grow completely at 37°C. For all suppression assays, plates were photographed every day for 4 days using Evolution MP Color Camera (Media Cybernetics) and Q Capture software. Images were imported into ImageJ software. Cells of interest were selected and *measure* tool was used to generate values for the integrated density, area and the mean signal intensity. Growth of a colony was quantified by measuring the signal intensity of the colony against the background using the formula, corrected signal intensity = Integrated Density – (Area of selected cell X Mean signal intensity of the background cell). Growth at 37°C was compared to growth at 30°C for each strain and then normalized to wild type. Quantitation was reported for the day where the temperature-sensitive mutant containing *STE11-4* plasmid showed around 60% of growth compared to the wild type, which was the maximum growth detected across various mutant combinations and independent trials. For most analysis, only the first or the second dilution of cells was used for the quantitation. All growth assays were performed in triplicate. Error bars show standard error of mean among the 3 trials.

### Quantitative PCR Analysis

Cells for qPCR were concentrated (OD A_600_ = 20) and spotted in 10 μl aliquots onto YEP-GAL (2% agar) for 24 h. Cells were spotted in six colonies per plate equidistant to each other and the plate center. All six colonies were harvested for each trial, and two separate trials were compared for each strain. The entire colony surface was scraped into 500 μl of distilled water, harvested by centrifugation, washed, and stored at −80°C. RNA was harvested by hot acid phenol chloroform extraction as described (Adhikari and Cullen, 2014). Samples were further purified using Qiagen RNeasy Mini Kit (Cat. 74104). RNA concentration and purity was measured using NanoDrop (NanoDrop 2000C). RNA stability was determined by 1% agarose Tris-Borate-EDTA (TBE, 89 mM Tris base, 89 mM Boric acid, 2 mM EDTA) gel electrophoresis.

cDNA libraries from RNA samples were generated using iScript Reverse Transcriptase Supermix (BioRad, 1708840). qPCR was performed using iTaq Universal SYBR Green Supermix (BioRad, 1725120) on BioRad (CFX384 Real-Time System). Fold changes in expression were determined by calculating ΔΔCt using *ACT1* mRNA as the housekeeping gene for each sample. Experiments were performed with biological replicates, and the average of multiple independent experiments were recorded. Primers for pPCR for *GIC2* were forward 5’-GCGCCAACAAGACAAATCACAAA-3’ and reverse 5’-GCAATTGCTCATCTTGGAATCC-3’. Primers for *GIC1* were forward 5’-GCCGAACAAGAACAACATCAA-3’ and reverse 5’-GTTTTGGCAGACCCATGTCTC-3’. Primers for *ACT1* were forward 5’-GGCTTCTTTGACTACCTTCCAACA-3’ and reverse 5’-GATGGACCACTTTCGTCGTATTC-3’ as published (Chavel *et al.*, 2010).

### DIC and Fluorescence Microscopy

Differential-interference-contrast (DIC) and fluorescence microscopy using FITC, CFP, YFP, Rhodamine, and DAPI filter sets were performed using an Axioplan 2 fluorescent microscope (Zeiss) with a Plan-Apochromat 100X/1.4 (oil) objective (N.A. 1.4) (cover slip 0.17) (Zeiss). Digital images were obtained with the Axiocam MRm camera (Zeiss) and Axiovision 4.4 software (Zeiss). Adjustments to brightness and contrast were made in Adobe Photoshop. Some images were obtained using structural deconvolution with a Zeiss Apotome filter. Multiple polarization events were assigned by examining cells over multiple focal planes by DIC and fluorescence microscopy. Time-lapse microscopy was performed on a Zeiss 710 confocal microscope equipped with a Plan-Apochromat 40x/1.4 Oil DIC M27 objective on a heated stage at 37°C. GFP was imaged using 488nm laser excitation and emission window from 496nm-572nm. Time-lapse Z stacks were captured at 10 min and 20 min intervals to monitor bud emergence and Cdc42p clustering.

### Budding Rate Assay

Cells were grown for 16 h on SD+AA medium. Using a toothpick, cells were removed from the plate, washed twice with water and resuspended in 1 ml water. 50 µl of cells were spread onto SD+AA or S-GAL+AA media and incubated at 30°C. Cells were visualized at 0, 5, 10 and 18 h intervals at 20X. Budding rate, ln(n) / ln(2), was determined as described (Hall *et al.*, 2014), where n is the number of daughters produced. Budding rate was adjusted for the time interval (5 h, 5 h and 8 h). More than 30 cells were examined for each interval.

### Time Lapse Microscopy

For time lapse microscopy, cells were placed onto semi-solid agarose pads that were prepared as described (Skinner *et al.*, 2013) with the following modifications. Approximately 700 µl of SD-URA media, prepared with agarose (1%), was placed inside 12 mm Nunc glass base dishes (150680, Thermo Scientific, Waltham, MA) and allowed to set at 25°C for 5 min.

For GFP-Cdc42p clustering in *cdc24-4* mutant combinations, cells were taken from colonies grown at 30°C for 16 h on SD-URA+G418 semi-sold agar media [2% agarose, 0.67% YNB without ammonium sulfate, 0.1% monosodium glutamate (MSG), 2% dextrose, 1 X amino acid stock without uracil, 0.36 mg/ml G418], resuspended in 7 µl of synthetic broth (0.67% YNB without ammonium sulfate, 0.1% MSG, 2% dextrose), and placed under the agarose pads by gently lifting the pad with a scalpel. 100 µl of water was placed in the dish adjacent to the agarose pad to prevent moisture loss, and the petri dishes were incubated at 37°C for 4 h. For co-localization of Sho1p-GFP and Gic2p-PBD-tdTomato, cells were prepared as above except imaged at 30°C. For GFP-Cdc42p clustering in cells carrying *MSB2*^Δ100-818^, 500 µl of saturated culture was washed twice with water and resuspended in 500 µl water. 5 µl was placed under the agarose pad, and the petri dish was incubated at 30°C for 10 h before imaging. For strains expressing GFP-Cdc42p and Cdc3p-mCherry, cells were prepared from colonies grown at 30°C for 16 h on SD-URA.

### FLIM-FRET Analysis

Plasmids containing the Cdc42p biosensor have been described (Smith *et al.*, 2013) and were a generous gift from Dr. Rong Li (Johns Hopkins University, Baltimore MD). Cells containing pRS316-pACT1-eGFP-CRIBCLA4-Cdc42p^ΔCAAX^–mCherry-CAAX (PC7137), and Q61L (PC7138) or D57Y (PC7139) derivatives, were grown on SD-URA medium to maintain selection for the plasmids. Semi-solid agarose pads were prepared as described above. Cells were taken from colonies grown for 16 h on semi-solid agar medium (SD-URA), resuspended in 7 µl of SD broth, and placed under the agarose pads.

Confocal Microscopy and FLIM was performed at SUNY-Buffalo North Campus Confocal Imaging Facility. FLIM-FRET has been described (Gratton *et al.*, 2003; Periasamy and Clegg, 2009; Osterlund *et al.*, 2015; Padilla-Parra *et al.*, 2015; Sun and Periasamy, 2015; Bassard and Halkier, 2018). FLIM-FRET images were acquired with a Simple Tau TCSPC 150 and HPM-100-40 GaAsP detector (Becker & Hickl GmbH) employing the direct coupled port of the LSM 710 and Zeiss Intune Laser excitation at 490 nm. At least 1000 photons per pixel were acquired for the highest count region for each cell.

FLIM-FRET data analysis was performed using SPCImage 7.3 (Becker & Hickl GmbH). Data was fit by a two-component exponential decay model and automatically generated Instrument Response Function (IRF). Bin factor, shift and offset were adjusted to obtain good fit (low χ^2^). For measuring the average lifetime, the decay curve was pooled for the region of interest. The mean fluorescence lifetime for two-component decay model was calculated according to the equation:

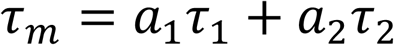

where *a_i_* and *τ_i_* are the amplitude and lifetime of the ith component. *τ_m_* is the average lifetime of the donor fluorescence. More than 20 cells were counted for each strain.

### Scanning Electron Microscopy

Scanning electron microscopy (SEM) was based on established methods (Piccirillo and Honigberg, 2011) and performed essentially as described (Basu *et al.*, 2016). For some experiments, cells were grown for 16 h in liquid media at 30°C. Cells were washed in 0.1 M sodium phosphate buffer pH 7.4 and diluted to about 10^6^ cells, which were passed over 0.2 micron Whatman nucleopore polycarbonate filter paper with a 10 mL syringe (GE Whatman, catalog #889-78084; BD Syringe, #309604). Cells were rinsed with one round of buffer by syringe, fixed with 1% glutaraldehyde for 15 minutes, and rinsed again. Cells were treated by a graded series of ethanol washes (30%, 50%, 70%, 85%, and 100%) by syringe to dehydrate the samples. The filter paper was removed from the holder, placed in a petri dish and treated with hydroxymethyldiazane (HMDS). Samples were placed at 4°C for 16 h and imaged the following day. All solutions were filter sterilized before use and stored in clean containers free of corrosion products. Samples were carbon coated and imaged on a FE-SEM (Field emission scanning electron microscope) Hitachi SU70.

### Image Analysis

Gic2p-PBD-tdTomato clustering was quantified as described in (Okada *et al.*, 2017) with following modifications. Raw fluorescence and DIC images were imported in ImageJ. DIC image was used to draw the cell boundary with the polygon tool. The same region of interest (ROI) was applied to the fluorescence image. Signal intensities for all pixels inside the ROI were exported as a CSV file using *save XY coordinates* option under *Analyze-tool* menu. A custom MATLAB (MATLAB R2016b, The MathWorks, Inc., Natick, MA) code (available on request) was designed to analyze the CSV files. In the analysis, mean and standard deviation (STD) of all the pixels of a cell were calculated. The pixel intensity that was greater than the mean+2 STD was selected. A region of the image where no cell was present was chosen as the background and the average pixel intensity of the background region was subtracted from the selected pixels of the cell. The signal intensity for each of these background subtracted pixels was normalized to the peak value which was set to 1. The sum of these normalized pixels was used to represent each cell. Over 60 cells were measured for each sample.

Time lapse microscopy images were processed in ImageJ. Grayscale fluorescence images were converted into maximum intensity projection and inverted. Quantitation of timing of Cdc42p clustering was performed as described (McClure *et al.*, 2016; Lai *et al.*, 2018) with following modifications. Threshold was applied to the 16-bit grayscale image to highlight cells and converted to binary image. Any merged cells were separated using watershed plugin after applying Gaussian Blur filter with sigma value of 2. Using *Analyze Particles* tab, the binary image was used to analyze the grayscale image to calculate mean intensity and standard deviation of each cell. Coefficient of Variance was then calculated from the mean and standard deviation and plotted as a function of time.

Kymographs were made on inverted maximum intensity projection image. Brightness and contrast was adjusted uniformly for each time frame and each strain. *Line* tool with pixel width of 1 was used to define region of interest and *Reslice* tool was used to generate slices for each time frame.

### Bioinformatics and Statistical Analysis

GO term analysis (Ashburner *et al.*, 2000) was performed using the Gene Ontology enRIchment analysis and visuaLizAtion algorithm (GOrilla) (Eden *et al.*, 2007; Eden *et al.*, 2009) using the two unranked lists mode. Genes encoding proteins associated with cell polarity were identified by GO term analysis [(Ashburner *et al.*, 2000), GO:0000282, cellular bud site selection; GO:0030010, establishment of cell polarity; GO:0031106, septin ring organization; GO:0006887, exocytosis; GO:0007120, axial cellular bud site selection; GO0007121, bipolar cellular bud site selection; GO:0030427, site of polarized growth]. GO terms and descriptions come from SGD (http://www.yeastgenome.org). Statistical significance for pair-wise comparisons was performed using Student’s t-Test in Microsoft Excel. For multiple comparisons, one-way ANOVA with Tukey test was performed in Minitab (www.minitab.com).

## ABBREVIATIONS

3-ATA: 3-Amino-1,2,4-triazole
5-FOA: 5-fluoroorotic acid
CFP: cyan fluorescent protein
D: dextrose
GAL: galactose
GAP: GTPase activating protein
GEF: guanine nucleotide exchange factor
GTPase: guanine nucleotide triphosphatase
GFP: green fluorescent protein
GLU: glucose
HA: hemaglutinin
HOG: high osmolarity glycerol response
Lat-A: Latrunculin-A
MAPK: mitogen activated protein kinase
MSG: Monosodium Glutamate
PAK: p21 activated kinase
qPCR: quantitative polymerase chain reaction
PM: plasma membrane
Rho: Ras homology
SDS-PAGE: sodium dodecyl sulfate-polyacrylamide gel electrophoresis
TBE: Tris-Borate-EDTA
TCA: trichloroacetic acid
WT: wild type
YFP: yellow fluorescent protein
YNB: Yeast Nitrogen base.

## ACKNOWLEDGEMENTS

Thanks to Matthias Peter (ETH Zürich), D. Lew (Duke University), John Pringle (Stanford University), Rong Li (Johns Hopkins University, Baltimore MD), Charlie Boone (University of Toronto), Alan Davidson (University of Toronto), Scott Emr (Cornell University), Peter Novick (Yale University), Erfei Bi (UPENN) and Mike Yu (SUNY-Buffalo) for providing strains and/or plasmids. Thanks to Steve Free (SUNY-Buffalo) for comments. Karnesh Jain (University of North Dakota) wrote the MATLAB code for quantifying Gic2p-PBD-tdTomato signal. Nadia Vadiae, Pu Zheng, Minakshi Mukherjee, Matt Vandermeulen and Shally Lin helped with experiments. Attendance at the 18^th^ annual workshop on FLIM and FRET microscopy at W.M. Keck Center for Cellular Imaging at the University of Virginia (hosted by Ammasi Periasamy), came from a travel award from the Histochemical Society (for A.P.). The work was supported from a grant from the NIH (GM#098629).

