## Supplemental Files for "Regulation of Bud Emergence by a MAPK Pathway"

#### SUPPLEMENTAL FIGURE LEGENDS

**Figure S1. Effect of fMAPK pathway on growth defect of the *cdc24-4* mutant.** (**A**) Typical example of cells of the indicated strains by DIC at 100X. Bar, 10 microns. Numbers refer to surface area of cells measured in Image J. Error shows the standard error of mean for three separate trials. 50 cells were counted in each trial. The differences in the values compared to wild type are significant. p-value < 0.01. (**B**) Immunoblot analysis. p44/42 recognizes P~ERK, including P~Kss1p and P~Fus3p as indicated. Pgk1p control for protein levels. Numbers refer to the ratio of P~Kss1p to Pgk1p relative to wild type. **(C)** Role of the fMAPK pathway (p*STE11-4*) on the growth of the *cdc24-4* and indicated mutant combinations. CTL, plasmid pRS316. See Fig.1B for details. (**D**) Volcano plot showing the distribution of fMAPK pathway target genes associated with cell polarity. Genes were graphed by $\log_2$fold change (X-axis, $|\log_2\text{fold change}| > 0.585$) and p-value (Y-axis, $< 10^{-5}$). GO terms are indicated by color. p-value <0.0001 for the genes shown. Asterisk denotes multiple GO term designations.

**Figure S2. *GIC2* is a target of the fMAPK pathway that is required for Cdc42p clustering and septin ring formation in the *cdc24-4* mutant. (A)** Quantitative PCR analysis showing relative fold changes in the mRNA levels of *GIC2* (left) and *GIC1* (right) in indicated strains. Error bars represent standard deviation among 2 replicates. Asterisk, p-value $< 0.05$. CTL, wild type. Other polarity targets genes were confirmed in [(Adhikari and Cullen, 2014) for *RSR1, BUD8,* and *RAX2*, and (Cullen *et al.*, 2004) for *MSB2*]. (**B**) Growth assay of wild type and *GIC1* mutant combinations at 37°C. CTL, plasmid pRS316. (**C**) Growth assay of the *cdc12-6* mutant harboring indicated plasmids at 30°C. Asterisk, p-value $<0.05$.

**Figure S3. Role of Gic2p in regulating filamentous growth. (A)** Invasive growth of the indicated strains by the PWA. Equal concentrations of cells were spotted onto YEPD media for 48 h. **(B)** Colony morphology for wild type, *gic1*Δ, *gic2*Δ and *ste12*Δ mutants. Equal concentrations of cells were spotted onto YEP-GAL medium for 24 h. Bar, 1 cm. **(C)** Cell morphology for wild type, *gic1*Δ, *gic2*Δ and *ste12*Δ mutants by the single cell invasive growth assay. Arrows indicate morphological defects. Bar, 10 microns. **(D)** Cdc12p-GFP localization in wild type, *ste12*Δ and *gic2*Δ mutants on SD and S-GAL media at 30°C. **(E)** Quantitation of septin localization. 50 cells were counted in each trial for three separate trials. Error bars represent standard error of mean. Asterisk, p-value $<0.05$. (**F**) The *gic1*Δ and *gic2*Δ mutant combinations (top) and Gic2p overexpression (bottom) along with controls were evaluated for the fMAPK pathway activity using *FUS1- HIS3* reporter assay (McCaffrey *et al.*, 1987). Cells were spotted in serial dilutions and incubated on selective media at 30°C for 2d.

**Figure S4. Effect of fMAPK pathway on growth defect of the *gic1Δ gic2Δ* mutant**. Role of hyperactive fMAPK pathway alleles on the growth defect of *gic1*Δ *gic2*Δ double mutant at 37°C alongside controls. Asterisk, p-value < 0.05. CTL, plasmid pRS316.

**Figure S5. Role of negative polarity complex on the activity of the fMAPK pathway and the mating pathway.** (**A**) The fMAPK pathway activity was compared among wild type and mutants in isogenic background using *FUS1- HIS3* reporter assay. Cells were spotted in serial dilutions and incubated on selective media at 30°C for 3d. (**B**) The *gps1*Δ mutant alongside controls was examined for invasive growth by the PWA (left) and single cell invasive growth assay (right). For the PWA, cells were spotted for 48 h on YEPD. The plate was photographed, then washed in a stream of water and photographed again. For the single cell assay, cells were spread onto S-Glu media for 16h. Bar, 10 microns. (**C**) At left, halo assay. Cells ($A_{600}$ 0.1) were spread on YEPD media and allowed to dry. 3 and 10 µl drops of 1mg/ml α-factor were applied to plates. Plates were incubated at 30°C and photographed at 24 h. The Bar, 1 cm. Representative image for 3 µl drop is shown. At right, halo diameters (cm) at 3 and 10 µl α-factor were measured by ImageJ. Error bars represent standard deviation among 3 independent trials. N.S., not significant.

**Figure S6. The fMAPK pathway mutant has growth defect under nutrient limiting conditions.** (**A**) Quantitation of the timing of GFP-Cdc42p clustering, Cdc3p-mCherry recruitment and bud emergence in wild-type cells (n=6) and *tec1*Δ cells (n=12) grown on semi-solid SD-URA media. Error bar represent standard error of mean. (**B**) Colocalization of Sho1p-GFP and Gic2p-PBD-tdTomato in wild-type cells grown in galactose for the indicated times at 15 min intervals. Red Arrow, bud emergence. Bar, 5 microns. (**C**) Same as B, except a representative example of wild type and *tec1*Δ mutant is shown. Black arrows, wandering Sho1p-GFP along the distal pole. (**D**) Growth assay of wild type and the *ste12*Δ mutant in glucose (GLU) and galactose (GAL). (**E**) Time series of single cell assay on wild-type cells and the *ste12*Δ mutant in GLU and GAL. Bar, 20 microns. (**F**) Quantification of budding rate of cells in panel E. Budding rate was calculated as described in Materials and Methods. Error bars, standard error of mean from 30 cells. Asterisk, p-value $<0.05$.

**Figure S7. Time lapse of bud emergence in fMAPK pathway mutants**. The following movies are also associated with the manuscript: (**A**) Time lapse of wild-type cells expressing GFP-Cdc42p and Cdc3p-mCherry in GAL. Time interval, 10 min. Time 0 is when septin hourglass splits into double ring marking cytokinesis. (**B**) Time lapse of *ste12*Δ mutant expressing GFP-Cdc42p and Cdc3p-mCherry in GAL. See panel A for details. (**C**) Second example of *ste12*Δ mutant expressing GFP-Cdc42p and Cdc3p-mCherry in GAL. Time 0 indicates start of experiment. (**D**) Third example of *ste12*Δ mutant showing merged DIC and Cdc3p-mCherry grown in GAL. See panel A for details.(**E**) Fourth example of *ste12*Δ mutant showing merged DIC and Cdc3p-mCherry grown in GAL. Time 0 indicates start of experiment. (**F**) Time lapse of *tec1*Δ mutant expressing GFP-Cdc42p and Cdc3p-mCherry in GAL. Time interval, 15 min. See panel A for details.

**Figure S8. Multiple growth sites by the fMAPK pathway.** (**A**) Merged image of DIC and rhodamine phalloidin staining of a cell expressing *MSB2*$^{\Delta 100\text{-}818}$. Serial sections in the plane of the Z-axis are shown. Bar, 5 microns. (**B**) At left, wild type and *MSB2*$^{\Delta 100\text{-}818}$ expressing Sec3p-GFP evaluated by single cell assay. At right, rhodamine phalloidin staining of a cell expressing Msb2p$^{\Delta 100\text{-}818}$.

**Figure S9. Time lapse of bud emergence during hyperactivation of the fMAPK pathway.** (**A**) Time lapse of wild-type cells expressing GFP-Cdc42p. Time interval, 10 min. (**B**) Time lapse of *MSB2*$^{\Delta 100\text{-}818}$ expressing GFP-Cdc42p. Time interval, 20 min.

**Figure S10**. **Role of overexpression genes in rescue of the growth defect of the *cdc24-4* mutant**. Wild-type cells and the *cdc24-4* mutant containing the indicated plasmids (Gelperin *et al.*, 2005) were spotted in serial dilutions onto SD-URA-LEU (GLU) and S-GAL-URA-LEU (GAL) media and incubated at 30°C for 2 d.

**SUPPLEMENTAL MOVIE LEGENDS**

**Movie 1.** Time lapse of wild-type cells expressing GFP-Cdc42p and Cdc3p-mCherry in GAL. Time interval, 10 min. Time 0 is when septin hourglass splits into double ring marking cytokinesis.

**Movie 2.** Time lapse of *ste12*Δ mutant expressing GFP-Cdc42p and Cdc3p-mCherry in GAL. See movie 1 for details.

**Movie 3.** Second example of *ste12*Δ mutant expressing GFP-Cdc42p and Cdc3p-mCherry in GAL. Time 0 indicates start of experiment.

**Movie 4.** Third example of *ste12*Δ mutant showing merged DIC and Cdc3p-mCherry grown in GAL. See movie 1 for details.

**Movie 5.** Fourth example of *ste12*Δ mutant showing merged DIC and Cdc3p-mCherry grown in GAL. Time 0 indicates start of experiment.

**Movie 6.** Time lapse of *tec1*Δ mutant expressing GFP-Cdc42p and Cdc3p-mCherry in GAL. Time interval, 15 min. See movie 1 for details.

**Movie 7.** Time lapse of wild-type cells expressing GFP-Cdc42p. Time interval, 10 min.

**Movie 8.** Time lapse of $\mathit{MSB2}^{\Delta 100\text{-}818}$ expressing GFP-Cdc42p. Time interval, 20 min.

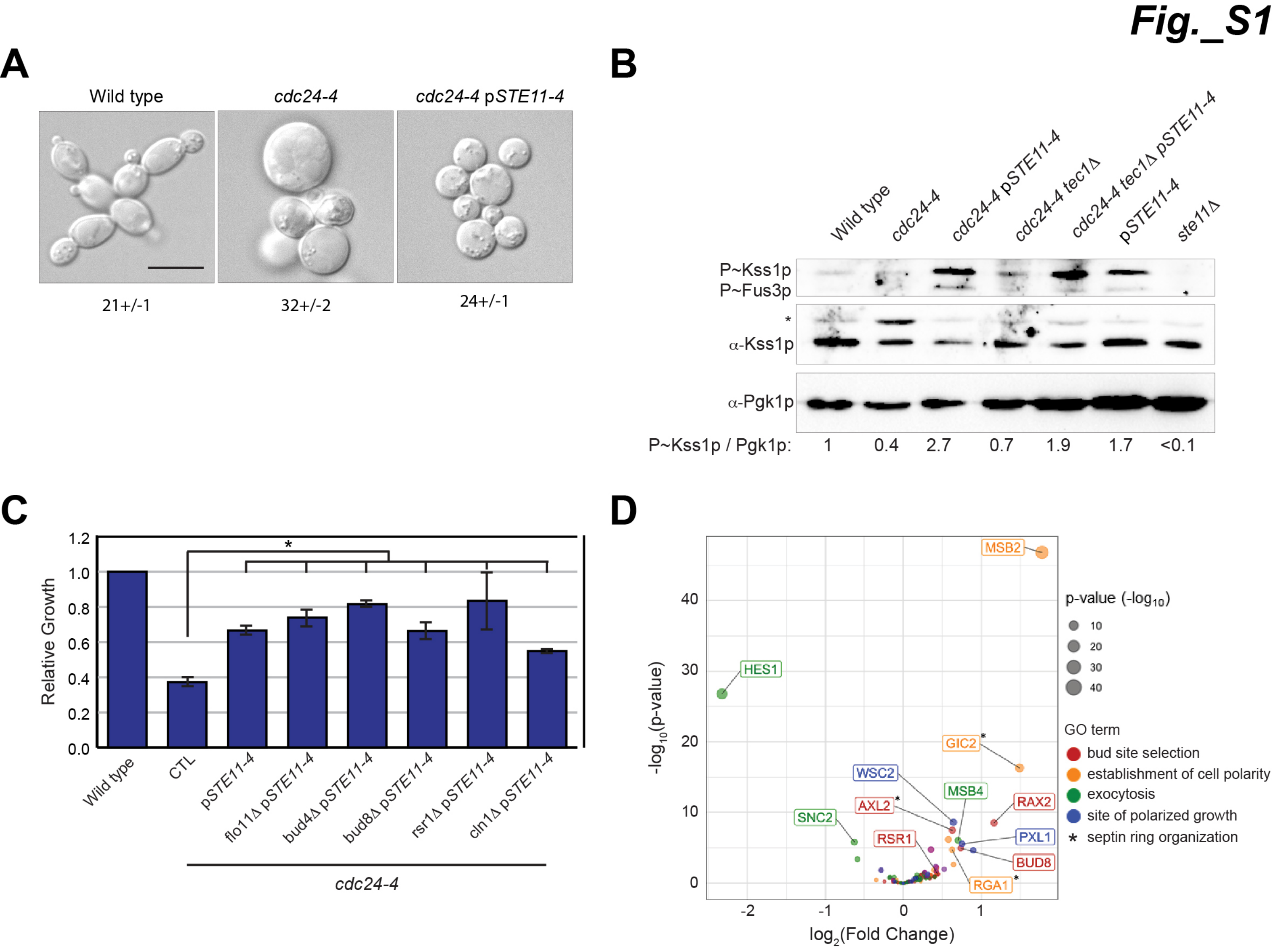
Fig._S1
A
Wild type
cdc24-4
cdc24-4 pSTE11-4
21+/-1
32+/-2
24+/-1
B
Wild type
cdc24-4
cdc24-4 pSTE11-4
cdc24-4 tec1Δ
cdc24-4 tec1Δ pSTE11-4
pSTE11-4
ste11Δ
P~Kss1p
P~Fus3p
*
α-Kss1p
α-Pgk1p
P~Kss1p / Pgk1p: 1 0.4 2.7 0.7 1.9 1.7 <0.1
C
Relative Growth
1.2
1.0
0.8
0.6
0.4
0.2
0.0
*
Wild type
CTL
pSTE11-4
flo11Δ pSTE11-4
bud4Δ pSTE11-4
bud8Δ pSTE11-4
rsr1Δ pSTE11-4
cln1Δ pSTE11-4
cdc24-4
D
MSB2
HES1
GIC2*
WSC2
MSB4
AXL2*
RAX2
SNC2
RSR1
PXL1
BUD8
RGA1*
-log10(p-value)
log2(Fold Change)
p-value (-log10)
10
20
30
40
GO term
bud site selection
establishment of cell polarity
exocytosis
site of polarized growth
* septin ring organization

***Fig._S2***

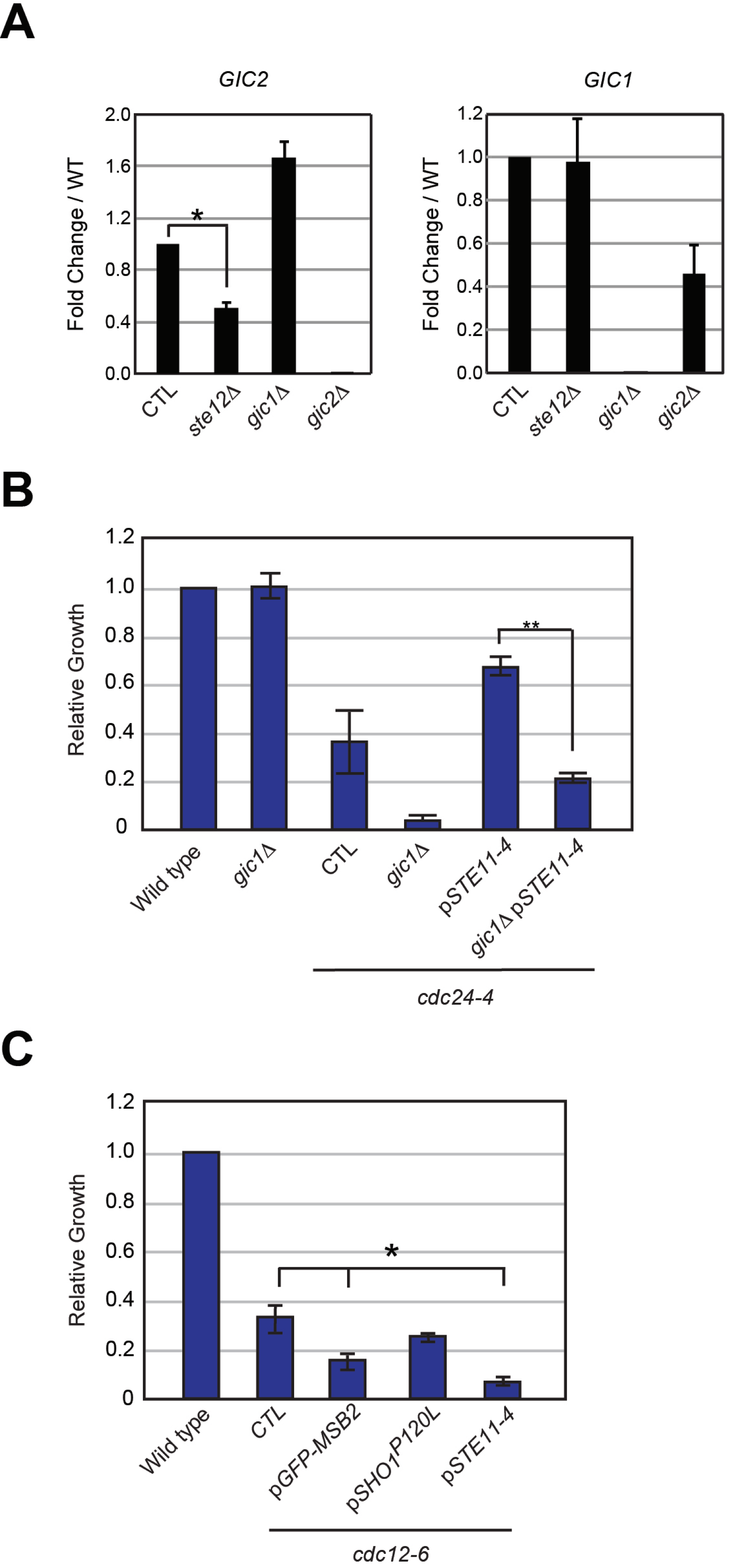

A
GIC2
Fold Change / WT
2.0
1.6
1.2
0.8
0.4
0.0
*
CTL
ste12Δ
gic1Δ
gic2Δ
GIC1
Fold Change / WT
1.2
1.0
0.8
0.6
0.4
0.2
0.0
CTL
ste12Δ
gic1Δ
gic2Δ
B
Relative Growth
1.2
1.0
0.8
0.6
0.4
0.2
0
**
Wild type
gic1Δ
CTL
gic1Δ
pSTE11-4
gic1Δ pSTE11-4
cdc24-4
C
Relative Growth
1.2
1.0
0.8
0.6
0.4
0.2
0
*
Wild type
CTL
pGFP-MSB2
pSHO1P120L
pSTE11-4
cdc12-6

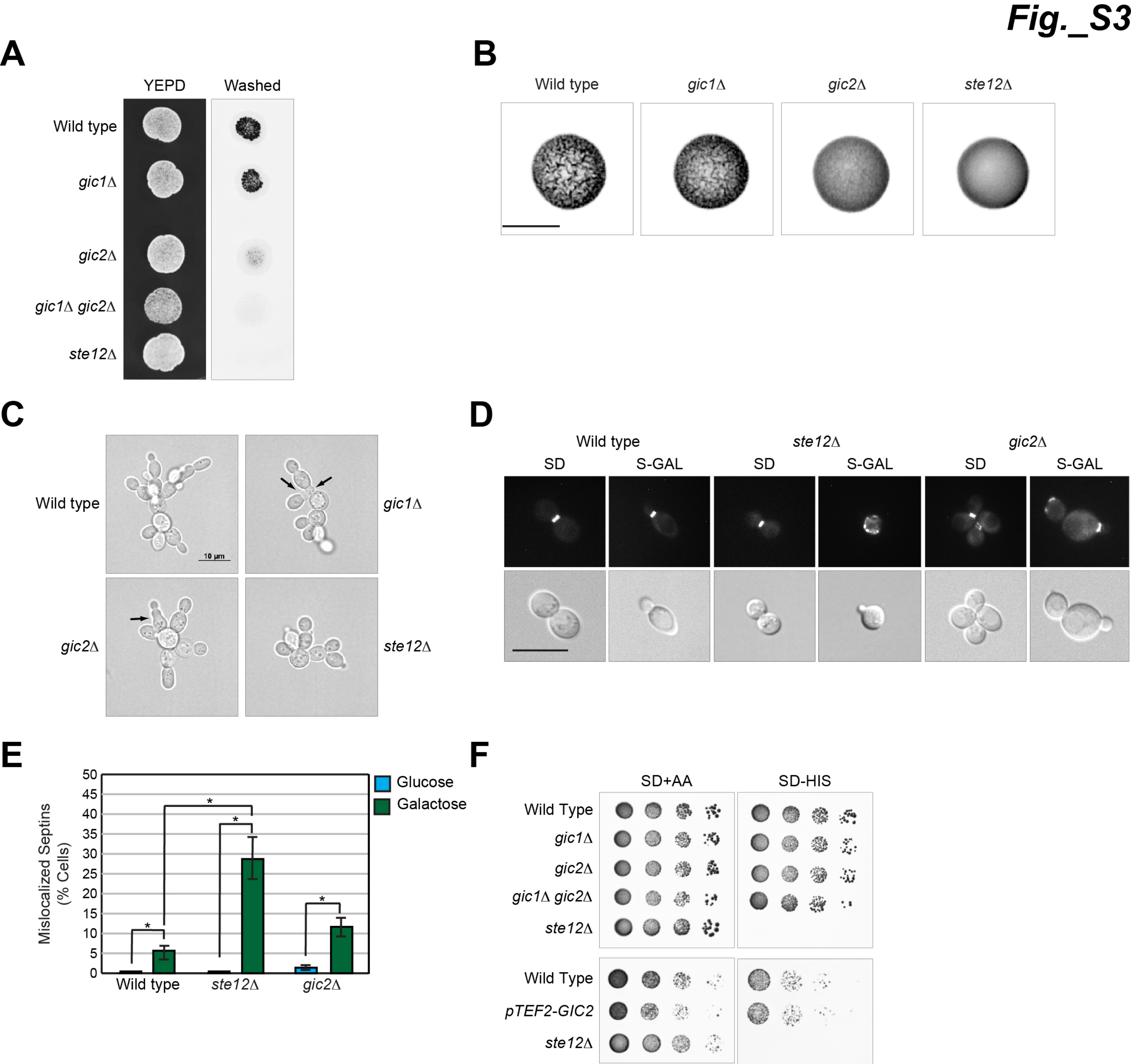
Fig._S3
A
YEPD
Washed
Wild type
gic1Δ
gic2Δ
gic1Δ gic2Δ
ste12Δ
B
Wild type
gic1Δ
gic2Δ
ste12Δ
C
Wild type
gic1Δ
gic2Δ
ste12Δ
10 µm
D
Wild type
ste12Δ
gic2Δ
SD
S-GAL
E
Mislocalized Septins (% Cells)
Glucose
Galactose
0
5
10
15
20
25
30
35
40
45
50
Wild type
ste12Δ
gic2Δ
*
F
SD+AA
SD-HIS
Wild Type
gic1Δ
gic2Δ
gic1Δ gic2Δ
ste12Δ
Wild Type
pTEF2-GIC2
ste12Δ

### Fig._S4

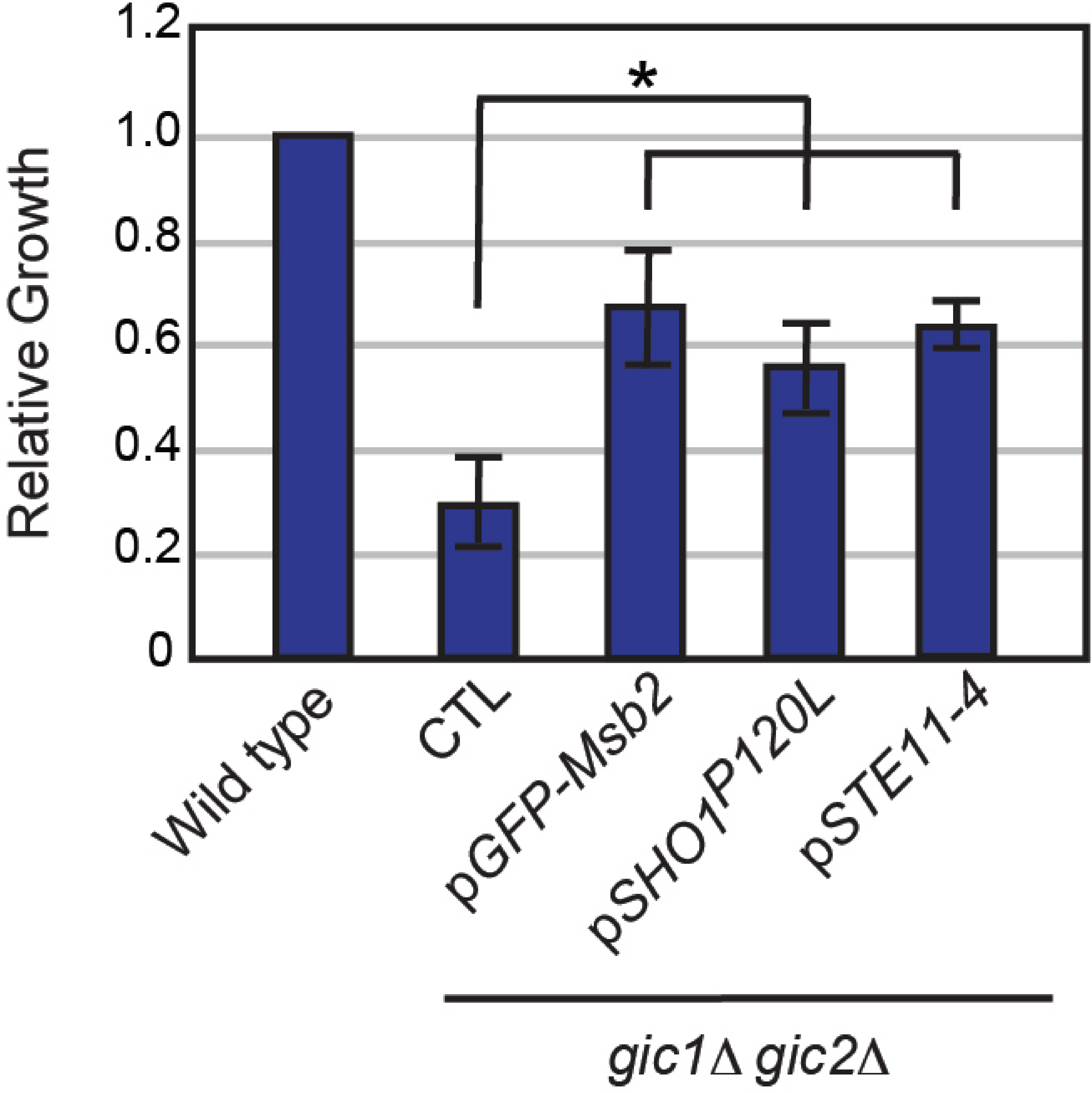
1.2
1.0
0.8
0.6
0.4
0.2
0
Relative Growth
*
Wild type
CTL
pGFP-Msb2
pSHO1P120L
pSTE11-4
gic1Δ gic2Δ

### Fig._S5

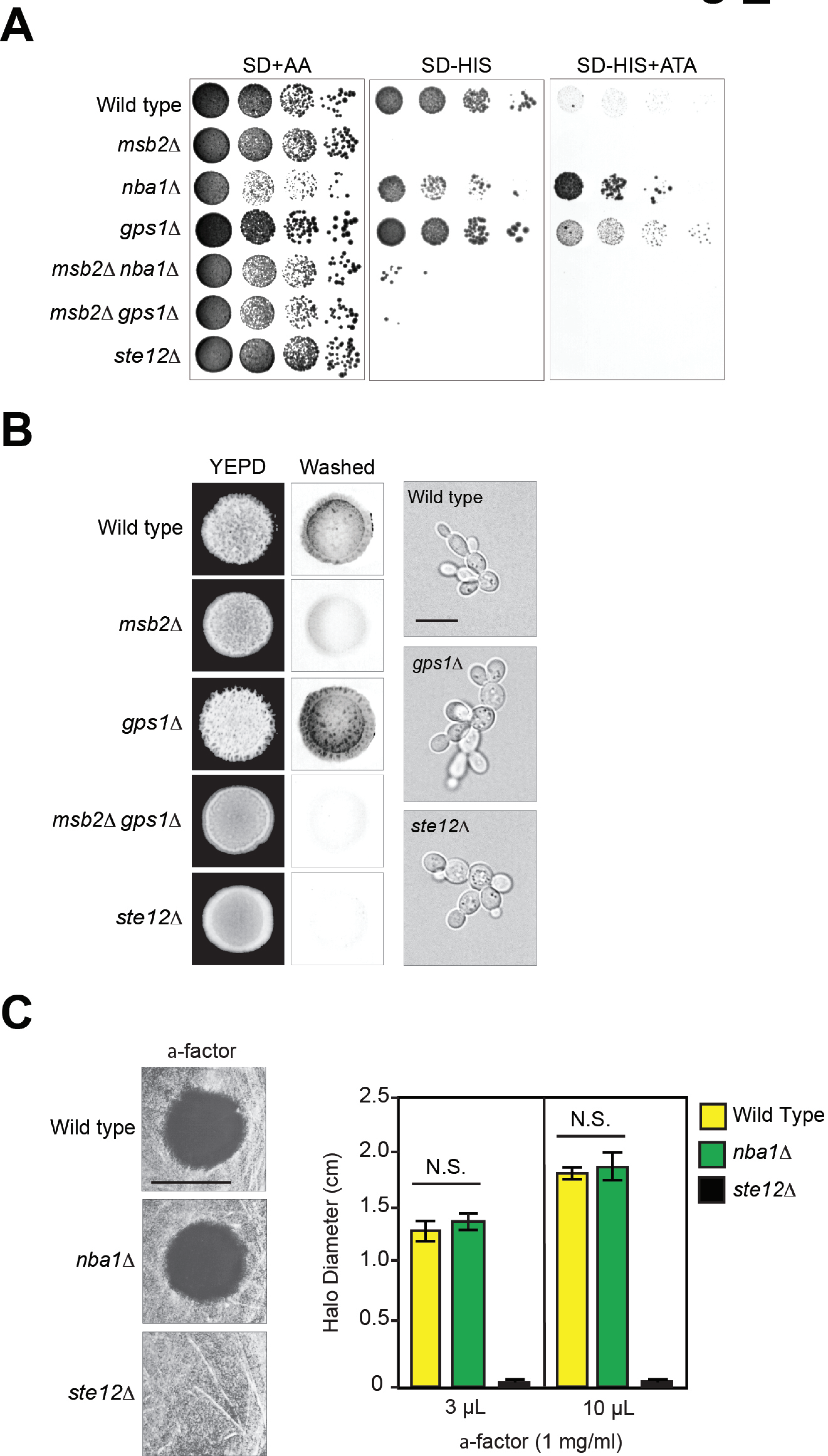
A
SD+AA
SD-HIS
SD-HIS+ATA
Wild type
msb2Δ
nba1Δ
gps1Δ
msb2Δ nba1Δ
msb2Δ gps1Δ
ste12Δ
B
YEPD
Washed
Wild type
msb2Δ
gps1Δ
msb2Δ gps1Δ
ste12Δ
Wild type
gps1Δ
ste12Δ
C
a-factor
Wild type
nba1Δ
ste12Δ
Halo Diameter (cm)
2.5
2.0
1.5
1.0
0.5
0
N.S.
N.S.
3 μL
10 μL
a-factor (1 mg/ml)
Wild Type
nba1Δ
ste12Δ

*Fig._S6*

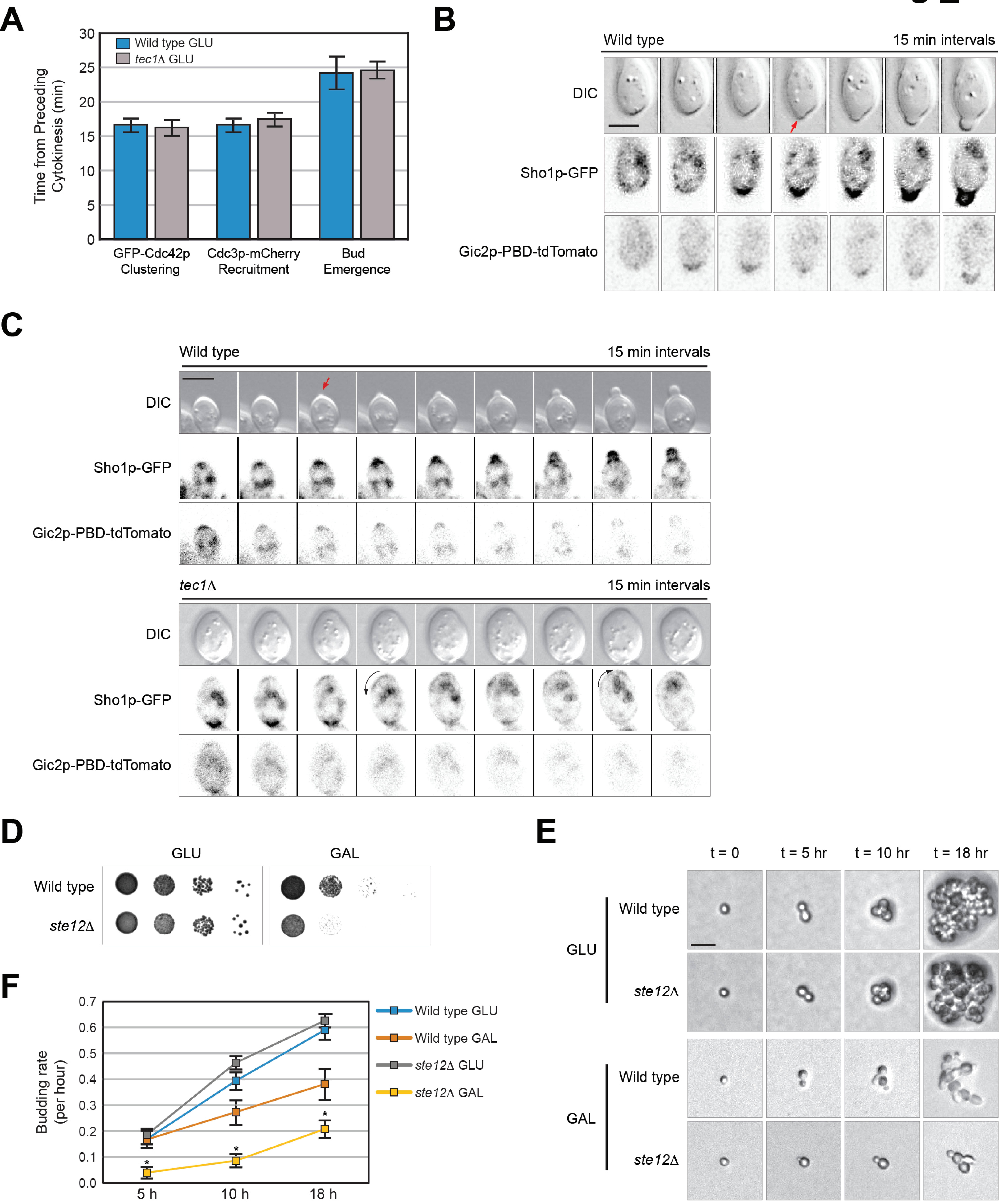

A
Wild type GLU
tec1Δ GLU
Time from Preceding Cytokinesis (min)
30
25
20
15
10
5
0
GFP-Cdc42p Clustering
Cdc3p-mCherry Recruitment
Bud Emergence
B
Wild type
15 min intervals
DIC
Sho1p-GFP
Gic2p-PBD-tdTomato
C
Wild type
15 min intervals
DIC
Sho1p-GFP
Gic2p-PBD-tdTomato
tec1Δ
15 min intervals
DIC
Sho1p-GFP
Gic2p-PBD-tdTomato
D
GLU
GAL
Wild type
ste12Δ
E
t = 0
t = 5 hr
t = 10 hr
t = 18 hr
GLU
Wild type
ste12Δ
GAL
Wild type
ste12Δ
F
Budding rate (per hour)
0.7
0.6
0.5
0.4
0.3
0.2
0.1
0.0
5 h
10 h
18 h
Wild type GLU
Wild type GAL
ste12Δ GLU
ste12Δ GAL

Fig._S7

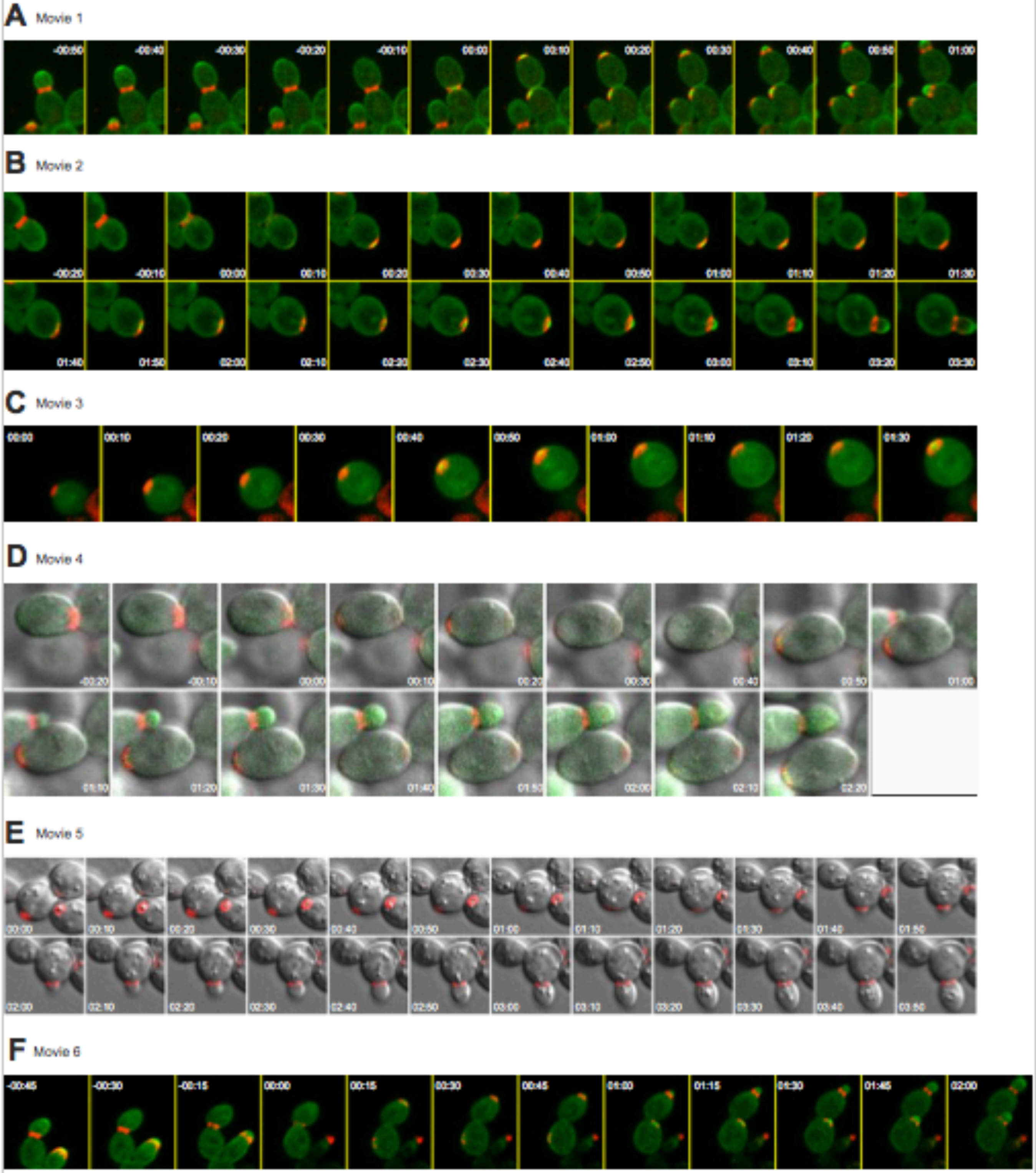

### Fig._S8

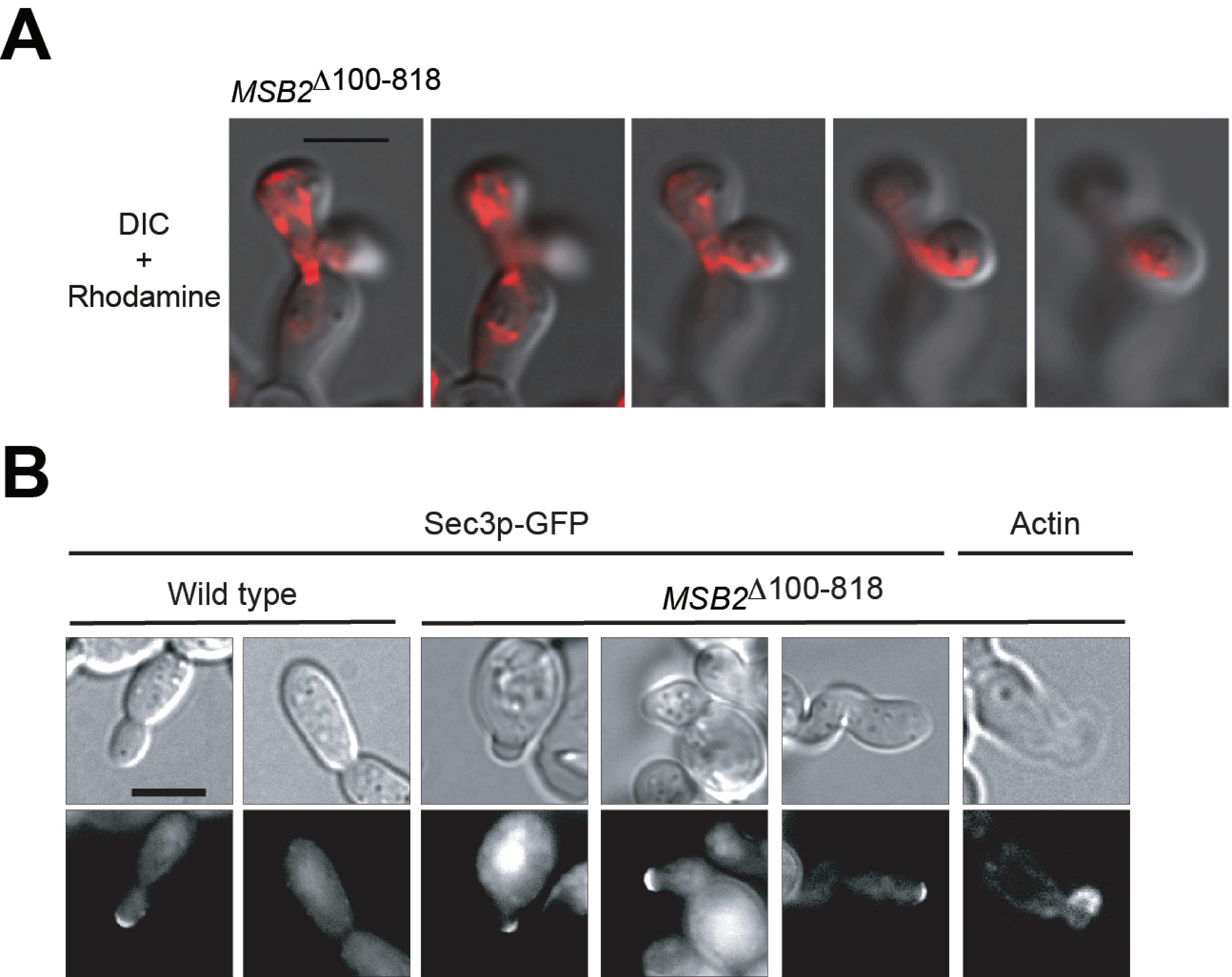
A
$MSB2^{\Delta 100\text{-}818}$
DIC
+
Rhodamine
B
Sec3p-GFP
Actin
Wild type
$MSB2^{\Delta 100\text{-}818}$

Fig._S9

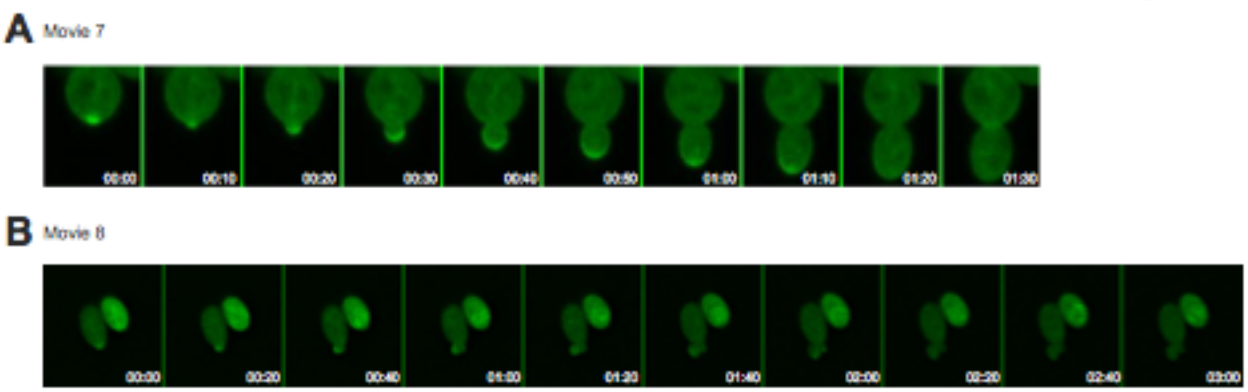
A
Movie 7
00:00
00:10
00:20
00:30
00:40
00:50
01:00
01:10
01:20
01:30
B
Movie 8
00:00
00:20
00:40
01:00
01:20
01:40
02:00
02:20
02:40
03:00
